# Inference of Synaptic Connectivity and External Variability in Neural Microcircuits

**DOI:** 10.1101/650069

**Authors:** Cody Baker, Emmanouil Froudarakis, Dimitri Yatsenko, Andreas S. Tolias, Robert Rosenbaum

## Abstract

A major goal in neuroscience is to estimate neural connectivity from large scale extracellular recordings of neural activity *in vivo*. This is challenging in part because any such activity is modulated by the unmeasured external synaptic input to the network, known as the common input problem. Many different measures of functional connectivity have been proposed in the literature, but their direct relationship to synaptic connectivity is often assumed or ignored. For *in vivo* data, measurements of this relationship would require a knowledge of ground truth connectivity, which is nearly always unavailable. Instead, many studies use *in silico* simulations as benchmarks for investigation, but such approaches necessarily rely upon a variety of simplifying assumptions about the simulated network and can depend on numerous simulation parameters. We combine neuronal network simulations, mathematical analysis, and calcium imaging data to address the question of when and how functional connectivity, synaptic connectivity, and latent external input variability can be untangled. We show numerically and analytically that, even though the precision matrix of recorded spiking activity does not uniquely determine synaptic connectivity, it is often closely related to synaptic connectivity in practice under various network models. This relation becomes more pronounced when the spatial structure of neuronal variability is considered jointly with precision.

## 1 Introduction

Modern interest in connectivity inference in neuroscience is quite broad in scope, ranging in scale from the microscopic properties of dendritic arbors to macroscopic cooperation across whole brain regions (Magrans de Abril et al, 2018). Even at a single scale, there are at least two distinct types of “connectivity” that are explored: functional connectivity and actual synaptic connectivity.

Many studies focus on inferring functional connectivity, which can broadly be defined as any statistical measurement of the functional interaction between neurons or other units in a neural system. A widely used method to infer functional connectivity at the level of local neural circuitry is to fit recorded neural activity to a generalized linear point-process model (GLM) which incorporates non-linearities when estimating effective coupling (Paninski, 2004; Pillow et al, 2008). These non-linearities correct the neuron’s responses to relate more to direct interaction and thus pertain more strongly to the underlying structure (Mishchencko et al, 2007). The accuracy of inference for GLMs has been evaluated *in silico* using simulations of non-linear Hawkes process models (Pernice et al, 2011). These models are idealized for GLMs as they correspond exactly to the statistical assumptions of the GLM inference algorithms so accurate inference of model parameters is to be expected. But the Hawkes model itself lacks the biophysical details accounted for in more mechanistic models such as networks of Hodgkin-Huxley (HH) style or Integrate-and-Fire (IF) neuron models. As such, functional connectivity inferred by GLMs applied to Hawkes process models is often interpreted not to approximate actual synaptic connectivity, but rather to represent the “effective” interaction between neurons with respect to the model network (Feldt et al, 2011; Poli et al, 2016)

Large-scale inference of synaptic connectivity between neuron pairs can be reliably performed using slice reconstruction or genetic mosaic analysis (Chiang et al, 2011), but such reliable approaches are lacking for *in vivo* applications. Less invasive extracellular recordings using large-scale calcium imaging or micro-electrode arrays that can be performed relatively safely *in vivo* but do not provide direct information about synaptic connectivity. Instead, the underlying connectivity structure of the recorded circuit influences the recorded activity.

Previous work has evaluated the relationship between functional and synaptic connectivity when GLMs are fit to spiking data subsampled from simulations of networks of leaky IF neurons (Lütcke et al, 2013; Zaytsev et al, 2015). Specifically, they assessed recovery of the ground truth structure from the *in silico* biophysical model against inferred coupling in the statistical model, but found relatively low accuracy of recovery overall.

Several other studies have proposed various methods for inferring synaptic connectivity, but since ground truth connectivity is not typically known for *in vivo* recordings, the accuracy of these methods has only been tested using *in silico* simulations. This approach is necessarily sensitive to parameter choices and underlying assumptions made in the design of the simulations. One common and important assumption in many such studies is a lack of correlated input from outside the recorded network (Kadirvelu et al, 2017; Mishchencko et al, 2007; Pernice and Rotter, 2013; Poli et al, 2016; Zaytsev et al, 2015), which is not a realistic assumption for *in vivo* recordings. Distinguishing the effects of this “latent” correlated input from direct connectivity – known as the “common input problem” – is notoriously difficult (Paninski, 2004; Pillow et al, 2008), but is necessary for accurate inference of connectivity from *in vivo* recordings.

Even when common input has been modeled in a network, it has typically been incorporated not through explicitly correlated external processes but rather via subsampling of the recurrent network (Brinkman et al, 2017; Lin et al, 2017; Lütcke et al, 2013), with the unobserved part inducing correlations only by way of the existing connections with the observed portion of the network. While this does indeed generate external correlations to the observed network, they have a different structure than correlations coming from a feedforward external layer projecting onto the entire recurrent population (Chambers et al, 2017).

An exact model of the relationship between connectivity and activity statistics in neural circuits is not known and most likely intractable, but there are simple mathematical expressions that provide accurate approximations to this relationship for various computational models (Baker et al, 2018; Krumin et al, 2010; Pernice et al, 2011; Trousdale et al, 2012) and these expressions can account for correlated external input. We evaluate how well synaptic connectivity can be inferred from estimates of spike train covariance under these approximations and how the quality of this inference depends on modeling assumptions and model parameters. We find that the precision matrix, *i.e* the inverse of the covariance matrix, of neurons’ spiking activity provides a good measure for inferring synaptic connectivity. We also find that inference can be greatly improved by accounting for the recorded neurons’ type (excitatory or inhibitory), tuning similarity, or distance, which are all quantities that can be measured or estimated during multicellular *in vivo* recordings. We test our conclusions using simulations of networks of adaptive exponential integrate-and-fire (AdEx) neuron models.

We begin by considering some simple motivating models and examples of functional measurements from them. Some of these models or measures are less practical for use in real data, and we will discuss their drawbacks in detail. We will then proceed to provide analytical details regarding the quality of network recovery based on functional measurements of spiking activity aggregated over large time windows. We then gradually introduce further biophysically realistic features into the model, and examine how the subsequent inference quality can be reduced or improved based upon knowledge (or lack thereof) of these covariates. Finally, we present a mean-field method for inferring properties of external latent variability for a neural circuit in mouse visual cortex.

## 2 Lessons on inferring connectivity from a simple stochastic rate model

As a motivating example, we begin by considering a simple, linear dynamical model (Dayan and Abbott, 2001) in which synaptic connectivity can be derived directly from observations of neuronal activity. The model is defined by

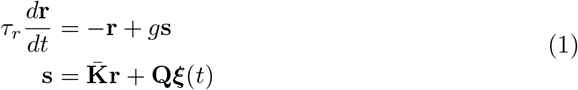

where **r**_*α*_(*t*) models the response of neuron *α* =1, 2,…, *N* as a low-pass filtered spike train or time-dependent firing rate, **s**_*α*_(*t*) models the synaptic input to neuron *α*, 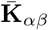 is the synaptic connection strength from neuron *β* to neuron *α, τ_r_* > 0 is a neural time constant, *g* > 0 is the neurons’ gain, ***ξ***(*t*) is *N*-dimensional standard Gaussian white noise modeling intrinsic noise and external synaptic input from outside the recurrent network, and **QQ**^T^ is the covariance matrix of the noise. This model defines a multi-variate Ornstein-Uhlenbeck (OU) process whenever 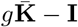 has eigenvalues with strictly negative real part (with **I** the identity matrix), which we assume to be the case. The stationary mean is

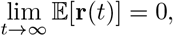

so **r**(*t*) should be interpreted as a mean-subtracted measure of firing rate. Correlations between neurons’ activity across time can be measured by the cross-covariance matrix

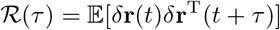

where 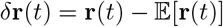, expectation is taken in the stationary state *t* → ∞, and **r**^T^ is the transpose of **r**. We wish to understand how the connectivity, 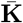, can be inferred from 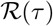. It can be shown that whenever 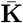 is normal, *i.e*. 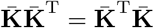, and **Q** = *ρ***I** is a multiple of the identity (implying that neurons receive independent external input), the off-diagonal entries of 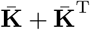 satisfy

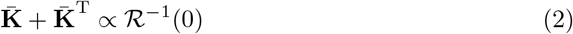

where 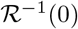 is the matrix-inverse of the zero-lag covariance, known as the *precision matrix* (see Appendix 13.2 for proof). The diagonal entries of 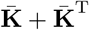 can be similarly derived, but we ignore them here because we are interested in connections between neurons and all of our network models lack self-connections. This can be seen as a generalization of the theory of Gaussian Graphical Models (GGM) wherein if 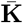 is symmetric with normally distributed non-zero elements, the exact precision perfectly encodes the conditional interactions between the neurons. However, connectivity in biological neuronal networks is not symmetric (**K** ≠ **K**^T^) and neurons are likely to receive correlated external input (**Q** not diagonal). Therefore, the functional connectivity inferred by the direct application of GGM methods to neural data does not necessarily correspond closely to synaptic connectivity. We further discuss the issue of statistical sampling of the inverse covariance in Section 10.

Fortunately, a regression theorem for OU processes (Gardiner, 2009) yields a more general expression for the off-diagonal entries of 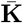 that is valid even when 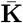 is not normal and **QQ**^T^ is not diagonal,

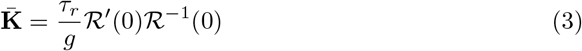

where 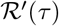 is the derivative of 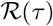 with respect to *τ*. Indeed, this estimator of 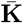 is analogous to estimates derived by expectation-maximization and maximum aposteriori (MAP) methods for multivariate AR(1) processes (Bishop, 2007; Singh et al, 2017), which are discrete-time analogues to OU processes. The derivative form in (3) is also analogous to differential covariance 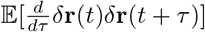 which has also been extended to the multivariate AR(2) process (Lin et al, 2017). Interestingly, this expression for 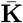 does not depend on **Q** at all, so it is not affected by correlated external input. Hence, using Eq. (3), synaptic connectivity can in principle be inferred directly from estimates of the cross-covariance between neurons’ activity under the model from Eq. (1). However, this approach has some critical shortcomings.

First, the model defined by Eq. (1) ignores the timescales of synaptic filtering, neuronal filtering, and external input variability that exist in biological neuronal networks. More specifically,

1. The model assumes that neural activity is transferred instantaneously to synaptic input, 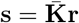, which ignores the temporal filtering imposed by synaptic kinetics.
2. The model assumes that neural activity is proportional to synaptic input, **r** = *g***s**, which ignores the temporal filtering imposed by neural membrane dynamics.
3. The model represents external input as Gaussian white noise, whereas external input to biological neuronal networks comes from the spiking activity of pre-synaptic neural populations, which is correlated across time.

The accuracy of Eq. (3) depends sensitively on these assumptions because the independence of Eq. (3) on **Q** relies on the fact that the contribution of **Q** to 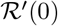 and 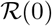 is the same, so **Q** cancels out in Eq. (3), but the same is not true of the contribution of 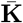. This difference is due to the timescales over which **Q** and 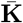 affect **r**.

Secondly, note that Eq. (3) requires evaluating 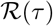 at small values of *τ*. This is problematic because fine-timescale dynamics are exactly what the model gets wrong (as outlined above), but also because large-scale multicellular recordings – such as those obtained from calcium imaging – often have low temporal resolution (though finer timescale dynamics can be inferred by deconvolution methods (Friedrich et al, 2017; Pnevmatikakis et al, 2017)). This makes it difficult to obtain accurate estimates of 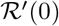 and 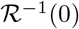 from data. Even in electrophysiological recordings that have fine temporal resolution, spike train correlations are often quantified from spike counts over long time windows (~250ms) due in part to the inherently low signal-to-noise ratio of spike train data (Cohen and Kohn, 2011).

We next consider a more general model that can capture arbitrary timescales of synaptic filtering, neuronal filtering, and external input correlation then consider inference methods that do not depend on these timescales.

## 3 Synaptic interactions cannot be computed from spike train covariability under a general linear model

We now consider a more general linear model of the form

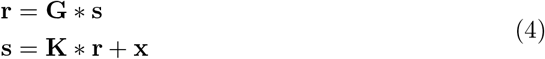

where * denotes matrix multiplication with each product replaced by a convolution over time (Trousdale et al, 2012),

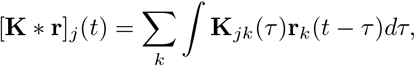

and where **x**(*t*) is some stochastic process modeling synaptic input from outside the local network. The synaptic connectivity kernel, **K**(*τ*), is an *N* × *N* matrix that accounts for synaptic weights as well as the time-course of synaptic filtering. The *N* × *N* diagonal matrix, **G**(*τ*), accounts for the filtering imposed by neural transfer of synaptic currents, **s**(*t*), to neural activity, **r**(*t*). This model resolves issues 1–3 mentioned above by accounting for arbitrary timescales of synaptic filtering, neuronal filtering, and external input noise.

To recover the OU process model in Eq. (1) from the more general model in Eq. (4), take

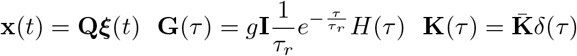

where *H*(*τ*) is the Heaviside function and *δ*(*τ*) is the Dirac delta function.

The second moments of this model over any timescale are determined completely by the cross-spectral matrix, defined as the Fourier transform of the cross-covariance matrix,

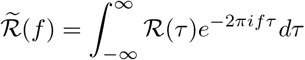

which can be written in closed form as

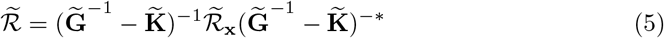

where ·^−1^ is the matrix inverse and ·^−^* is the inverse of the conjugate-transpose, 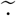 is the Fourier transform, and we have omitted the explicit dependence of 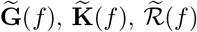, and 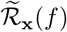 on frequency, *f*, for notational convenience. The *N* × *N* matrix, 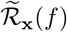, is the cross-spectral matrix of **x**(*t*), defined as the Fourier transform of the cross-covariance matrix, 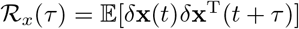.

From Eq. (5), it can be seen that the same cross-spectral matrix, 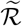, can be produced by two different connectivity matrices, 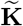, together with different matrices, 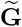 and 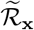. Hence, without knowledge of 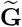 and 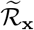 or additional assumptions on model parameters, one cannot infer 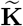 directly from measurements 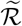. In practice, one does not typically have knowledge of pairwise external input correlations or neural response properties to constrain 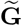 and 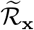 in neural recordings.

Furthermore, 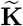 cannot be derived from 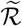 even when 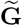 and 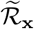 are known. To see this, consider the case where 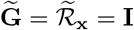 (corresponding to **G**(*τ*) = **I***δ*(*τ*) and **x**(*t*) = ***ξ***(*t*)) and note that derivation of 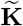 is equivalent to derivation of 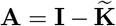. But **Σ** = **A**^−1^**A**^−^* is invariant to unitary transformations of **A**, *i.e*., to multiplication of **A** by a matrix, **U**, satisfying **UU**\* = **I**. Hence, multiple connectivity matrices, 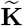, produce the same correlation structure, 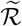, even when 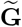 and 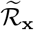 are fixed. This was pointed out in previous work (Pernice and Rotter, 2013), which assumed diagonal 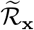 and inferred 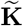 under an assumption of sparsity. However, high quality inference of 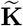 in that study was only possible when sparsity was lower than that observed in local cortical circuits (Jiang et al, 2016; Levy and Reyes, 2012) and, perhaps more importantly, the assumption of diagonal 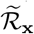 – which implies uncorrelated external input – is not justified in cortical populations.

Note that 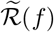 uniquely determines 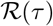 and other measures of spike train correlation such as spike count covariance and spike count correlation. Therefore, since 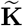 cannot be derived exactly from 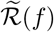, it cannot be derived from any of these other measures of spike train covariability either.

Eq. (5) was derived for the linear model in Eq. (4), which is arguably more biologically realistic than the model in Eq. (1), but is still a gross simplification of real neural circuit dynamics. Specifically, the model in Eq. (4) does not account for the nonlinearity of neural transfer. However, Eq. (5) has been shown to provide an accurate approximation for more biologically realistic networks of spiking neuron models (Baker et al, 2018; Trousdale et al, 2012) and non-linear Hawkes process models (Krumin et al, 2010; Pernice et al, 2011). Hence, we conclude that, under a wide class of models, synaptic interactions cannot be derived explicitly in terms of spike train covariance in the presence of unknown external input covariance. This is an example of the common input problem (Soudry et al, 2013) under which common or correlated input to neurons cannot be distinguished from direct synaptic connectivity between them.

A precise derivation of synaptic connection strengths in terms of spike train covariance is therefore perhaps too ambitious of a goal. Below, we weaken this goal to argue that, in practice for networks of randomly connected neurons, Eq. (5) allows us to infer the presence or absence of synaptic interactions between pairs of neurons with a great deal of accuracy.

## 4 A simpler goal: inferring undirected sparsity structure from precision

Instead of trying to infer the entire connectivity kernel, **K**(*τ*) or 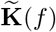, we aim only to infer its sparsity structure, *i.e*., which neuron pairs are connected. First note that the zero-frequency connectivity kernel,

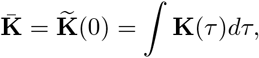

represents the matrix of total synaptic strengths. We then decompose 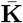 as

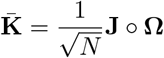

where ∘ is the element-wise (Hadamard) matrix product of synaptic weights, **J**, with a binary adjacency matrix, **Ω**. The 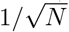 scaling permits stability of network dynamics when **Ω** and **J** are random matrices and promotes excitatory-inhibitory balance and asynchronous dynamics for large *N* (Renart et al, 2010; Van Vreeswijk and Sompolinsky, 1996; van Vreeswijk and Sompolinsky, 1998). There is evidence that synaptic weights in cultured populations of cortical neurons scale similarly (Barral and D’Reyes, 2016).

We then evaluate Eq. (5) at *f* = 0 and rescale all terms by 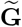 to obtain the simpler expression

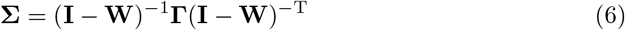

where 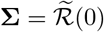 is the low-frequency covariance between neural activity,

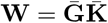

is the normalized synaptic weight matrix,

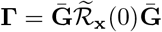

is the normalized external input covariance matrix, and 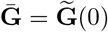 is the diagonal matrix of gains. Note that the inverse-conjugate-transpose, ·^−^*, in Eq. (5) is replaced by a inverse-transpose, ·^−T^, in Eq. (6) because the zero-frequency cross-spectral density between real-valued processes is real-valued (Yaglom, 1962). Note also that, since **Ḡ** is diagonal, **W** has the same sparsity structure as 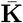, which is captured by **Ω**.

The matrices **Σ, Γ**, and **W** have natural interpretations. Since low-frequency susceptibility, **G**_*αα*_, represents the gain of neuron *α*, *i.e*., the derivative of the neuron’s f-I curve, **W**_*αβ*_ represents the connection strength from neuron *β* to neuron *α* scaled by the post-synaptic neuron’s sensitivity to inputs. Similarly, **Γ**_*αβ*_ represents the low-frequency covariance between external inputs to neurons α and *β* scaled by the sensitivity of both neurons.

Finally, **Σ**_*αβ*_ is proportional to the spike count covariance between neurons *α* and *β* over long time windows, which is a widely used measure of correlated variability (Cohen and Kohn, 2011; Doiron et al, 2016). It is also proportional to the low-frequency covariance between the neurons’ firing rate fluctuations. This means that **Σ** can be estimated using low temporal resolution measures of neural activity, such as those approximated by calcium imaging. Therefore, focusing on low-frequency interactions resolves the issue of temporal resolution described above.

Motivated by the theory established in Sections 1-3, we seek to infer connectivity using measurements of the low-frequency precision matrix,

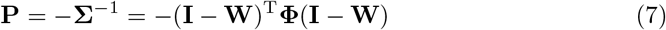

where **Φ** = **Γ**^−1^. Note **P** is distinct from the zero-lag temporal precision in Eq. (2) that is analogous to classical GGM theory. We will consider **P** our primary functional measure of interest throughout the remainder of this paper. Motivation for this choice comes from the fact that many functional measures of connectivity are functions of **P** (Kadirvelu et al, 2017; Lin et al, 2017; Pernice and Rotter, 2013; Poli et al, 2016; Yatsenko et al, 2015). In addition, for the classes of random networks we consider, the entries of **P** are approximately normally distributed for large network size (*N*), shown in Appendix 13.4, which makes inference easier to describe and understand. Also, **P** can be easily estimated from multicellular recordings by numerically inverting the sample spike count covariance matrix or low-frequency cross-spectral matrix between neurons’ activity. We discuss the estimation of **P** from data in more detail in Section 10 and in the Discussion. Until then, we evaluate the ability to infer connectivity from a perfect estimate of **P**.

Our main goal is then to infer the matrix of undirected binary connections, **Ω** + **Ω**^T^, from knowledge of precision, **P**, under the assumption that Eq. (7) is satisfied and that **Ω** is the binary adjacency matrix for **W**. We do not enter into this problem with expectations of fully recovering **Ω** + **Ω**^T^ for any network, but rather we seek to understand the underlying factors that contribute to a high degree of association between **Ω** + **Ω**^T^ and **P** under different network models.

Most literature on inferring connectivity in neuronal networks has focused on the simple case of uncorrelated external input (**Γ** and **Φ** diagonal) (Kadirvelu et al, 2017; Mishchencko et al, 2007; Pernice and Rotter, 2013; Poli et al, 2016; Zaytsev et al, 2015) and we will initially follow suit by assuming **Γ** ∝ **I**. We will later relax this assumption. In this case Eq. (7) reduces to

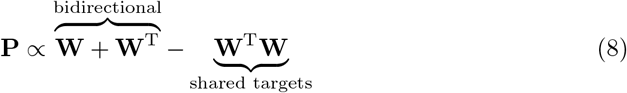

for the off-diagonal elements. The first term represents bidirectional connectivity in that it is non-zero at an entry only if there is a connection between the corresponding neurons, in at least one direction, *i.e*., only if **Ω** + **Ω**^T^ is non-zero at that entry. An entry of the second term is non-zero whenever the corresponding neurons share some post-synaptic targets. More generally, this term is larger in magnitude when the two neurons share more post-synaptic targets. Knowledge of the first term would give perfect inference of bidirectional connectivity, so the second term can be considered a source of noise when trying to infer **Ω** + **Ω**^T^ from **P**. A main intuition from Eq. (8) is the roughly linear relationship between **P** and **W** + **W**^T^, as demonstrated in Fig. 1a,b with similar results previously observed in studies of general linear point-process models (GLMs) (Mishchencko et al, 2007). This relationship is construed by the error term which arises from shared post-synaptic targets.

**Figure 1.**
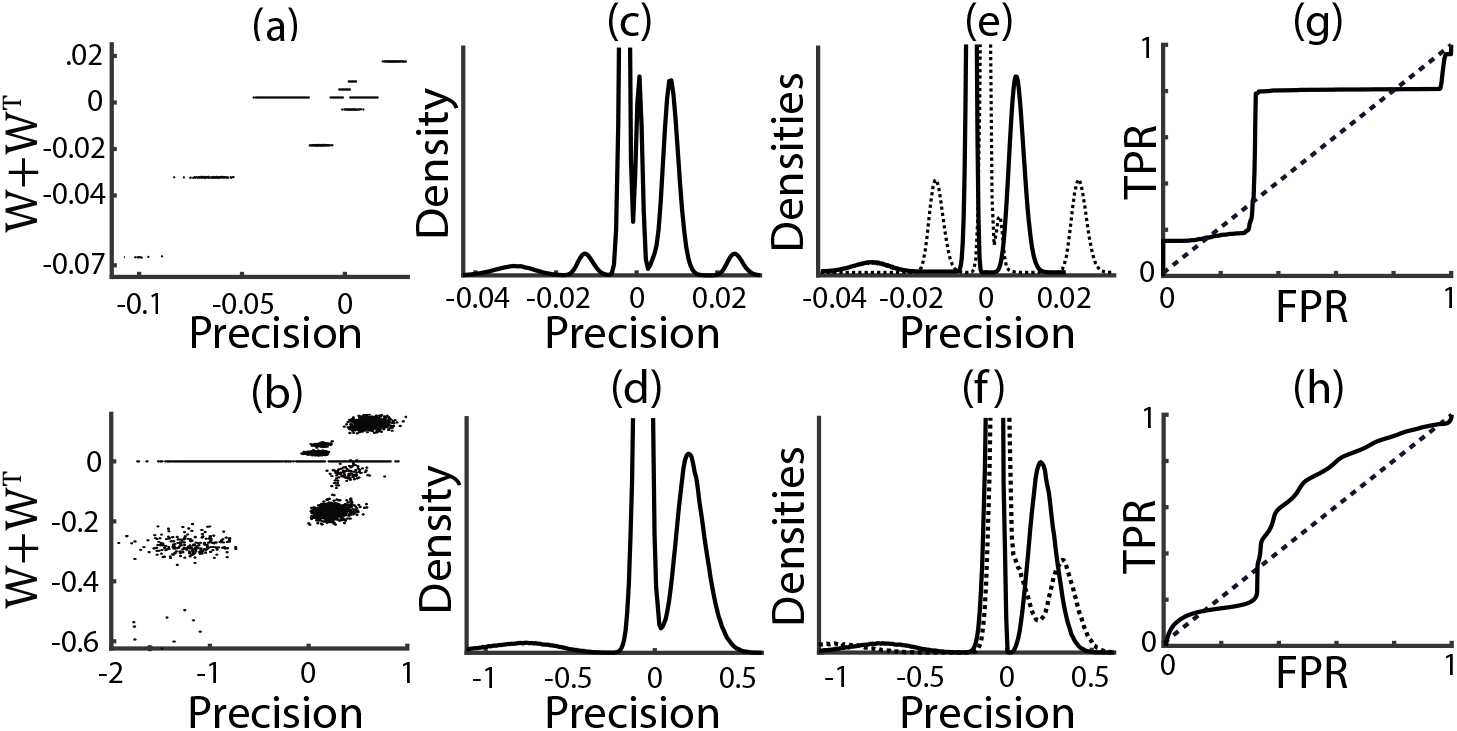
Inference in a simple model under a simple partition scheme results in low AUROC. (a) Scatter plot of 3 × 10^4^ randomly sampled pairwise values from a randomly generated precision structure of *N* = 2000 neurons versus the corresponding values of pairwise bidirectional connectivity. (c) Total empirical density of the values in (a). (e) Empirical densities of the values in (a) partitioned into connected (dotted line) and unconnected (solid) pairs; the connected distribution is no longer down-weighted by the low probability of connection. (g) ROC curve for distinguishing connected versus unconnected pairs in (e), with the dashed line as reference to a random classifier. (b,d,f,h) Same as (a,c,e,g) but in a higher noise, stronger strength regime. AUROCs: (g) 0.6084, (h) 0.5904.

We test this relationship by generating **W** according to various random graph models, initially with gains fixed at unity (**G** = **I**) for simplicity. All of the network models we consider contain *N_e_* excitatory (e) and *N_i_* inhibitory (i) neurons and obey Dale’s law (with *N* = *N_e_* + *N_i_*, *q_e_* = *N_e_*/*N* = 4/5, and *q_i_* = *N_i_*/*N* = 1/5). The connectivity matrix can be decomposed into four blocks where 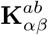 denotes connections from neuron *β* in population *b* = *e, i* to neuron *α* in population *a* = *e, i*. We start with a simple block-wise Erdos-Renyi model with normally distributed synaptic weights defined by

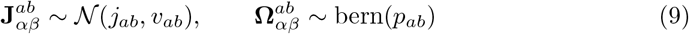

where all random variables are assumed to be independent. Hence, *j_ab_* denotes the mean synaptic strength, *v_ab_* the variance of synaptic strengths, and *p_ab_* the connection probability from population *b* = *e, i* to population *a* = *e, i*. We enforce Dale’s Law on the synaptic strengths by truncating the normal distribution onto the corresponding half-intervals of ℝ^+^ for excitatory and ℝ^−^ for inhibitory neuron types, but this truncation has a small effect when 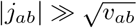.

To quantify the performance of network recovery, we utilize Receiver Operator Characteristic (ROC) curves, which are a common and reliable metric. ROC curves are generated by taking some set of values referred to as the *score* and assigning positive and negative classes by comparing the values against some threshold, and then counting the true and false positive rates (TPR/FPR) as the threshold itself varies to span the set of scores. In our case, the values in the precision matrix serve as the scores and the classes are initially partitioned into the simple connected versus unconnected sets. It is important to note that in the context of network recovery, the aforementioned model details combine to generate an approximate mixture distribution on the precision values. A randomly chosen value in the precision matrix takes the form

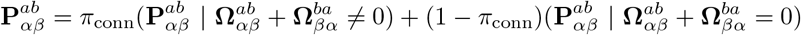

where *π_conn_* is a binary variable corresponding to whether the randomly chosen pair is connected. This mixture model form helps to further justify the appropriateness of the ROC metric.

We now perform an initial analysis of structural recovery using randomly generated networks following two basic regimes under which the coefficient of variation of **J** 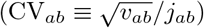 is near-zero or near-one, which we will refer to as the low-noise and high-noise regimes. The scatter plots in Fig. 1a,b reflect this escalation in noise, which ultimately causes the multi-modal mixture distribution structure apparent in Fig. 1c,e to collapse to two or fewer observable modes in Fig. 1d,f. The multi-modal shape of these distributions of precision values are due to the mixing of excitatory and inhibitory neurons as well as uni- and bi-directionally coupled motifs, all of which are later considered as additional information used by other partition schemes in Section 5.

The ROC curves for this initial partitioning are are shown in Fig. 1g,h. As observed, recovery of network structure is quite poor under this setting, yielding area under the ROC curve (AUROC) of around 0.6 in both cases. Note that all discussion and use of AUROC throughout this paper is in a folded sense, that is AUROC ∈ [0.5, 1] where any AUROC which would originally return a value in [0, 0.5) is folded back into the rightward interval. This convention accounts for situations in which a given method or measure is interpreted more appropriately as an anti-classifier.

Figs. 1g,h demonstrate that inferring connectivity by thresholding all pairs of precision values simultaneously yields poor recovery of synaptic connectivity. Fig. 1e,f demonstrate why this occurs: either there is a great deal of total overlap between the connected (dotted) and unconnected (solid) distributions, or the unconnected sub-groups are alternately dispersed between the peaks of the connected density. We next show that inference of connectivity from precision can be improved by conditioning on cell type.

## 5 Using cell-type labels can improve inference of connectivity

Above, we showed that thresholding precision, **P**, can give poor inference of connectivity (Fig. 1d,h). However, this conclusion was reached under the assumption that we had no information about whether the recorded neurons were excitatory or inhibitory. Indeed, the multimodal densities of precision values (Fig. 1b,c,f,g) are partly due to their representing multiple pre- and post-synaptic cell types. We will now show that by conditioning on this additional information, we can vastly improve our quality of inference.

In neural recordings, estimates of cell type can often be obtained by genetic labeling or classification of spike waveforms. To illustrate how inference can be improved by accounting for contextual data such as cell type, we utilize several families of data *masks* analogous to those used in (Lin et al, 2017) to explicitly specify how the elements of the precision matrix may be partitioned based on conditional information. A family of masks *M*(*a, b*; *m*) parameterized either by sub-populations *a, b* and/or by connection type m defines the set of values from the precision matrix which correspond to motifs of the selected type. These sets give rise to distributions over their elements and, so utilization of different masks will always specify a different set of ROC curves, each illustrating differing levels of inference quality.

If cell and connection types are known, then the richest contextual set of masks is

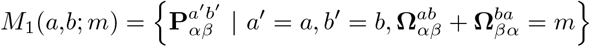

where *m* = 0 corresponds to the precision values between unconnected neurons, *m* = 1 corresponds to the precision values computed between pairs of uni-directionally coupled neurons, and *m* = 2 to the precision values between bidirectionally coupled neurons. Information regarding cell-type is granted through specification of sub-populations *a,b* ∈ {*e, i*}, which further restricts the conditional precision values to the block of the matrix which corresponds to that neuron type. For a given sub-population, inference using *M*_1_ allows the comparison of distributions *M*_1_ (*a, b*; 1) or *M*_1_ (*a, b*; 2) against the shared null (unconnected) group *M*_1_ (*a, b*; 0) within the ROC analysis. For example, the ROC curve computed by comparing the distribution of values in *M*_1_(*e, e*; 0) to those in *M*_1_(*e, e*; 1) (denoted *e* → *e* in figure legends) quantifies how well uni-directionally connected pairs of excitatory neurons can be distinguished from unconnected pairs of excitatory neurons when bi-directionally connected pairs have been removed and excitatory neurons are labeled.

Mask *M*_1_ is only applicable in situations where ground truth is available (since one needs to know which neurons are bi-versus uni-directionally coupled) and hence is generally only applicable to *in silico* network simulations. It is however useful for explaining the behavior of the other masks. A more reductive mask is then

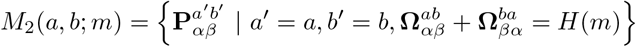

where *H*(·) is the Heaviside step function. As such, *m* in *M*_2_ may only take the values zero or one denoting the null (unconnected) and positive (connected) groups, thus causing the bidirectional motifs to be combined into the same set of values as the unidirectional. The null groups are still separated on a sub-population basis however, which is arguably the most important aspect. So mask *M*_2_ distinguishes only cell type, and combines the values across connected pairwise motifs. For example, the ROC curve omputed by comparing the distribution of values in *M*_1_(*e, e*; 0) to those in *M*_1_(*e, e*; 1) quantifies how well connected pairs of excitatory neurons can be distinguished from unconnected pairs when excitatory neurons are labeled. This corresponds to the situation faced when inferring connectivity from experimental recordings in which units are labeled by cell type.

Finally, the simplest mask is

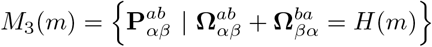

which is a common mask used on *in vivo* data (Vinci et al, 2018; Yatsenko et al, 2015) as well as *in silico* experiments following Dale’s Law (Chambers et al, 2017; Lin et al, 2017; Lütcke et al, 2013; Pernice and Rotter, 2013; Poli et al, 2016). Like mask *M*_2_, it only distinguishes null and positive connections, but now it does so without knowledge of sub-population membership. The ROC curve computed by comparing the distribution of values in *M*_3_(1) to those in *M*_3_(0) quantifies how well connected pairs can be distinguished from unconnected pairs when neurons have not been labeled by cell type, which is how the ROC curves in Fig. 1g,h were computed. In this case, there are three different null groups interspersed within the original eight positive (connected) classes from *M*_1_. It is this multitude of classes distinguished by *M*_1_ which are responsible for the multiple modes in the distributions from Fig. 1.

Utilizing mask *M*_1_ to distinguish between cell types and connection types provides a clear separation between the precision densities of each type (Fig. 2a-f, compare to Fig. 1a-f) and a dramatic improvement of inferred connectivity (Fig. 2g,h; compare to Fig. 1g,h). In the network with less synaptic variability, an AUROC of around 0.6 when using *M*_3_ was improved to multiple AUROC values all near 1 (near perfect classification) when using *M*_1_. For the network with greater synaptic variability, AUROC values were also generally improved by using *M*_1_ in place of *M*_3_ (see Fig. 2h caption).

**Figure 2.**
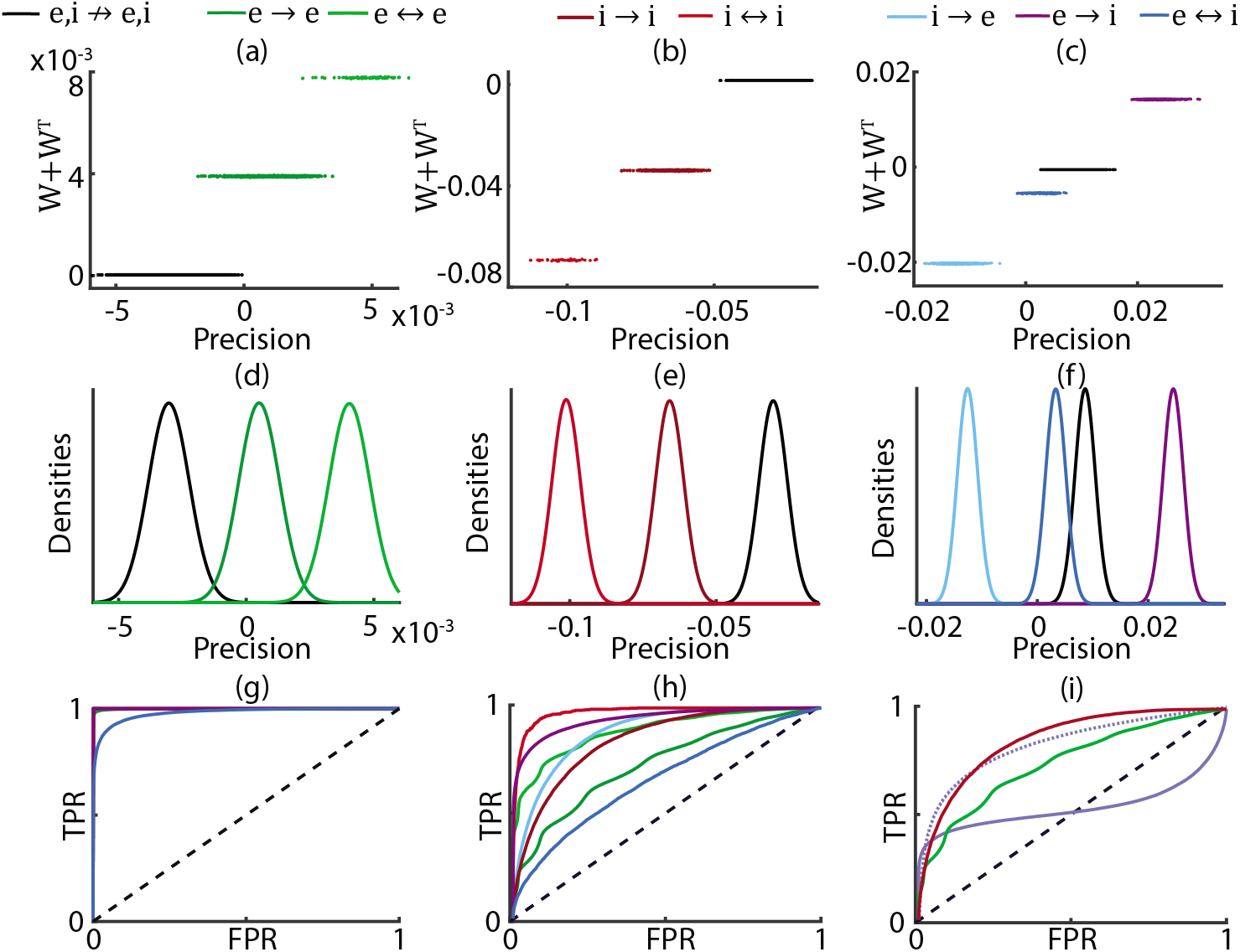
Utilizing more informative partition schemes (masks) improves inference quality. (a-c) Scatter plots of 26, 335 precision values versus corresponding bidirectional synaptic strength for the same structure used in Fig. 1, but now separated by excitatory/inhibitory subgroups (mask *M*_1_). Black always represents the null (unconnected) group for each subtype, but note this group is now different across subtypes whereas in Fig. 1 it was shared across all distributions. (d-f) Empirical densities of partitioned precision values in a-c. (g) ROC curves for distributions shown in d-f. (h) ROC plots of a similar network under a higher noise, stronger strength setting; same as values in Fig. 1b subject to mask *M*_1_. (i) ROC curves resulting from applying mask *M*_2_ to values in Fig. 1b (higher noise, stronger strength). Dotted line represents the absolute value of the centered cross-population. AUROCs two second decimal place, in order consistent with legend: (g) 1, 1, 1, 1, 1, 1, 0.98 (h) 0.73, 0.89, 0.85, 0.98, 0.95, 0.89, 0.66 (i) 0.74, 0.85, 0.54, as well as 0.83 for the dotted line. The networks are identical to Fig. 1.

In neural recordings, even if we know cell types, we typically do not know whether a particular pair value in precision corresponds to a bidirectional or unidirectional motif, so the application of mask M1 is not realistic for real neural data. The application of mask *M*_2_ in Fig. 2i represents the ROC curves that are produced in a more realistic setting in which recorded cells are labeled, but the nature of the connected motif is unknown. This is still a substantial improvement over the unlabeled data (Fig. 1e-h). It is important to note however that by combining the cross-population distributions (excitatory-inhibitory pairs; *a* = *e, b* = *i* or *vice versa*), there is substantial loss of inference in the strict sense because the null group is nested between the three connected groups. Such behavior is detectable as the ROC curve crossing the diagonal reference (Fig. 2i, solid purple curve), but this is easily corrected by taking the absolute value of the centered precision as the score to be thresholded (Fig. 2i, dotted curve).

In summary, accounting for additional information within the model, such as cell type labels or motif structures can improve the inference of synaptic connectivity.

## 6 An analytical expression for AUROC clarifies its dependence on parameters

This finding that the use of additional model information improves inference in simulations is encouraging, but we also wish to understand how sensitive inference quality can be as a function of the chosen parameter values. Thankfully, for the model we consider, this problem is analytically tractable and results in a direct function relating our model parameters to the area under the ROC curve. This function also directly reveals a number of qualitative features, many of which were previously discovered via *in silico* studies.

While there is some loss of information in reducing the full ROC curve to a single scalar value, the AUROC nonetheless provides a robust and widely used measure for quantifying the accuracy of recovery as parameters of the networks change.

The AUROC may be calculated analytically if applied to normally distributed scores (see Appendix 13.3 for a review of this theory). For the network types considered within this paper, the resulting precision values under mask *M*_1_ will be approximately normally distributed in the large network limit, a result proven in Appendix 13.4. We may thus explain the resultant AUROC for M_1_ as a bijective function of *discriminability D* (a.k.a, *sensitivity index, signal-to-noise ratio, Fisher’s criterion, Rayleigh’s quotient*)

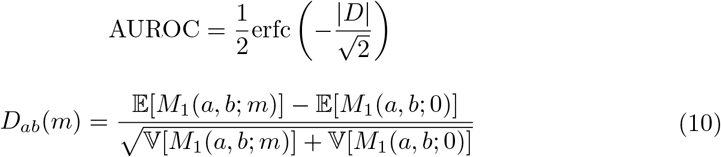

where erfc(·) is the complementary error function. We have defined our discriminability in terms of mask *M*_1_ whereby we distinguish unidirectional, bidirectional, and unconnected distributions from one another over all cell types. It may theoretically be possible to extend this theory to the other masks as well, however some difficulty which arises in this extension is the fact that the measures become more complicated mixtures of normal distributions for which similar expressions may not always be easily re-derived.

Under certain network architectures, we may describe *D* as a direct function of the underlying network parameters by evaluating the moments in Eq. (10) for the precision structure specified by Eq. (7).

Under the Erdos-Renyi assumptions of Eq. (9) together with the case of cell-type specific independent external white noise 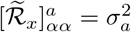 and randomly distributed inverse-gains with the first two moments parameterized as 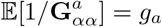 and 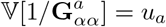, we derive the required quantities for *D* in Supplemental Section 3, giving

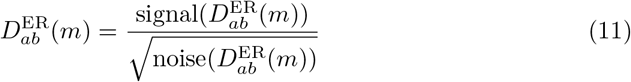

where

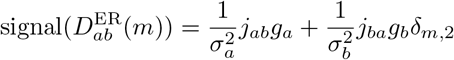

and

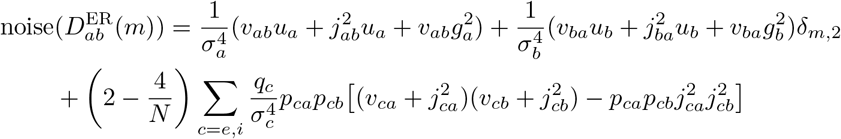

Here, *δ_m,n_* is the Kronecker delta. Some studies have numerically explored how the AUROC changes as functions of the parameter space: (Kadirvelu et al, 2017) showed how AUROC decreases for larger network sizes and (Pernice and Rotter, 2013) showed that it tends to increase for sparser networks. A direct analysis of Eq. (11) confirms these qualitative features and uncovers several dependencies of discriminability on various parameters, which we now review.

### AUROC is monotone decreasing in N

The dependence of 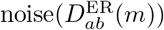 on network size, *N*, as well as the independence of 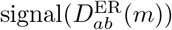 on *N* implies that discriminability will always be larger for smaller networks. One interpretation is that smaller network size in conjunction with the high sparsity levels (*p* ≪ 1) leads to fewer actual realizations of post-synaptic targets in the network, which forms the major component to the confounding variance across the precision distributions. For real neural networks however, *N* is likely to be quite large and so we will focus on the thermodynamic limit (*N* → ∞) of Eq. (11), which is accurate for even moderately large *N*.

### AUROC is monotone decreasing in the variance of synaptic weights

The dependence of 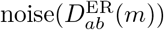 on synaptic weight variance, *v_ab_*, as well as the independence of 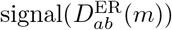 on *v_ab_* implies that increased variance of synaptic weights reduces discriminability. This is a very straightforward result pertaining to the prevalent modeling practice of having randomly distributed synaptic weights, with (Lütcke et al, 2013; Pernice and Rotter, 2013; Poli et al, 2016) being among the few studies that utilize fixed (zero variance) synaptic strengths.

### AUROC exhibits nontrivial dependence on the mean of synaptic weights

For non-random synaptic weights, the discriminability is monotone decreasing in *j_ab_*, implying that weaker synaptic strengths lead to better inference. Intuitively, this is due to the fact that as *j_ab_* → 0, the signal between the connected and null distributions goes to zero at a slower rate than the noise collapses. The situation becomes more complicated when synaptic weights are variable (*v_ab_* > 0), where discriminability now achieves a maximum at some particular value of *j_ab_* determined by the other parameters of the network (see Supplemental Section 4.1) and vanishes as *j_ab_* → 0 or ∞. Thus, variability of synaptic weights changes the qualitative dependence of discriminability on the mean synaptic weight, and in the presence of synaptic variability, there exists some level of synaptic strength ideal for inference.

### AUROC depends on neuron type

There are several items to note with respect to differences in discriminability over multiple neuron types. For even populations, that is when *a* = *b*, bidirectional connections will always be easier to distinguish than unidirectional connections from unconnected neurons (a very visible property in Fig. 2a-c). The opposite holds for the odd populations *a* ≠ *b* where bidirectional connections induce a “cancellation” at the level of the signals, due to the fact that *j_ie_* > 0 and *j_ei_* < 0.

### AUROC exhibits nontrivial dependence on network sparsity

It has been shown numerically (Pernice and Rotter, 2013) and follows from analysis of Eq. (11) that the AUROC → 1 as *p* → 0. More interesting behavior emerges for intermediate levels of sparsity, tending to dense networks. Supplemental Section 4.2 provides the analysis which leads to the following results. Under the reparameterized form of Eq. (11), the equation will have a minimum somewhere within the interval 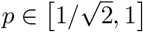 so long as the relative magnitude of synaptic variance is sufficiently weaker than the mean synaptic strengths. For settings involving larger variances, the discriminability will become monotone decreasing for denser levels of connectivity.

From these qualitative properties, or directly from Eq. (11), we may use our *a priori* knowledge to generate a random network that gives any pre-specified AUROC level. This makes it difficult to compare the effectiveness of various classification methods when each is taken over a different set of parameters. We cannot, for instance, directly compare and contrast the results in (Kadirvelu et al, 2017) and (Pernice and Rotter, 2013) as they possess different sets of network parameters. Furthermore, all observed AUROC’s from all purely *in silico* studies are a result of choices of numerous parameter values. This raises the concern that each individual method or measure may perform better or worse within different regions of the parameter space.

## 7 The effects of network structure on inference of connectivity

The Erdos-Renyi assumption of the previous section greatly simplifies the analysis, but real neural networks possess properties directly opposed to the strong assumptions of that model (Song et al, 2005). To account for some of these descrepencies, we will extend the analysis to frameworks that account for two cases: when there are correlations between the average strength and the out-degree of a neuron, and when the in- or out-degrees follow a heavy-tailed distribution. The results show that inference is made more difficult by including these more biophysically realistic features.

So far, we have only considered networks with a simple block Erdos-Renyi connectivity structure. We now consider two additional network models which incorporate more realistic features of neural networks. The first is the *copulated Erdos-Renyi* model (CER), motivated by some experimental results that suggest correlations between synaptic strength and connection probabilities (Jiang et al, 2016) and defined by

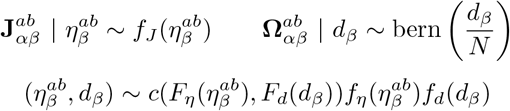

which obtains specifiable correlation levels *ρ* between average synaptic strengths and out-degrees for some some collection of arbitrary marginal densities *f* with cumulative distributions *F* bound together by some specified copula density *c*. For this model we do require that connection probabilities be homogeneous across the network (*p_ab_* = *p*) however the only other requirements we place on the general case involve the moment matching or parameterization of

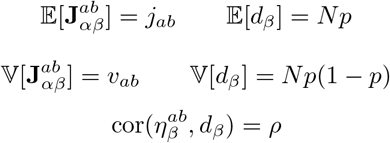

to induce consistent first and second order statistics with the simpler Erdos-Renyi model considered before. Though this general model could potentially apply to any dependence structure, for the purposes of our analysis we will assume Gaussian copulae and marginals for ease of analysis.

The previous discriminability analysis may be repeated for this model, revealing the following modification to Eq. (11)

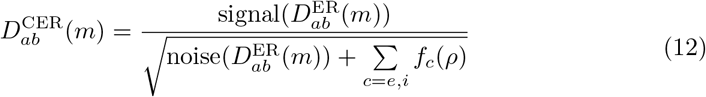

where the signal and noise are the same as those in Eq. (11). The form of the sub-population dependent function *f_c_*(*ρ*) in terms of the copulated correlation *ρ* is derived in Supplemental Section 3.1, and is rather large and complicated but is positive for our parameters and thus acts as an additional decrement to discriminability. Hence, correlations between neurons’ out-degrees and their synaptic weights can make accurate inference of synaptic connectivity more difficult. This point is demonstrated numerically in Fig. 3 (compare red to blue).

**Figure 3.**
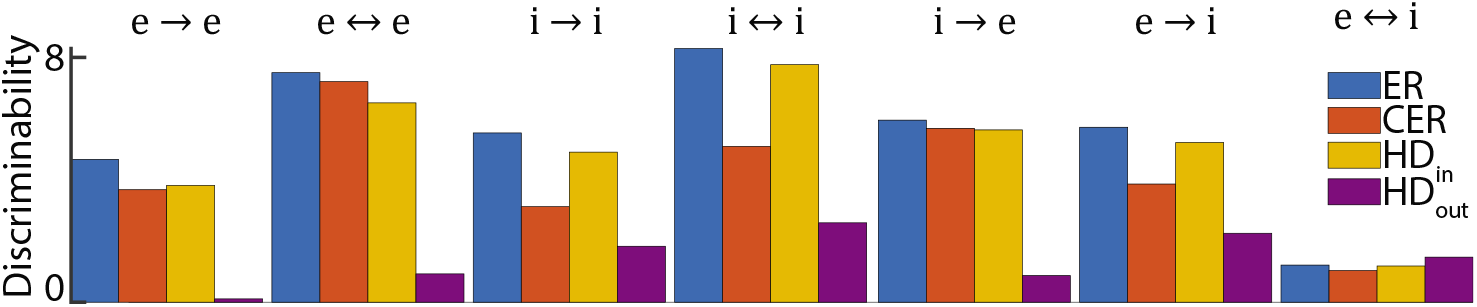
Mean discriminability for ten randomly generated precision structures of *N* = 2000 neurons over the four network types, partitioned over all subgroups. The slight increase in the bidirectional cross-population (*e* ↔ *i*) for HD_out_ is due primarily to the non-Gaussian nature of the distribution.

Yet another source of potential additional variability comes from the assumed form of the degree distribution, which in the Erdos-Renyi case is binomial. An existing generalization of Erdos-Renyi connectivity to allow specifiable heterogeneous in- or out-degree (HD_in/out_) distributions is used, most notably to include a power law in the form of a generalized Pareto distribution (Pyle and Rosenbaum, 2016)

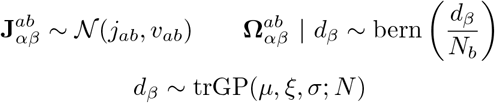

where trGP(⋯; *N*) denotes the Generalized Pareto distribution truncated at a maximum value equal to the number of neurons in the network. We will always hold *μ* and *ξ* fixed then numerically estimate the value of *σ* which induces a mean network density equivalent to the ER model (*N_p_*). Note that without truncation this value would be *σ* = (*N_p_* − *μ*)(1 − *ξ*), but the truncation shifts the optimal value in a non-linear fashion.

Extending the discriminability analysis to this model is only possible for Pareto distributions of in-degrees as the out-degree model does not admit a Gaussian central limit as disccused in Appendix 13.4. The in-degree case yields

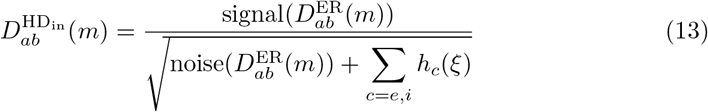

where *h_c_*(*ξ*) is the positive hyperparameter function from the Pareto variability. As with Eq. (12), the additional terms are positive and thus reduce discriminability. Hence, Pareto-distributed in-degrees can make inference of synaptic connectivity more difficult, as compared to an Erdos-Renyi model. This point is demonstrated numerically in Fig. 3 (compare yellow to blue). While our analysis does not extend naturally to Pareto-distributed out-degrees, a numerical comparison of this case shows reduced discriminability Fig. 3 (compare purple to blue).

In conclusion, network structure can affect discriminability and, specifically, different deviations of connectivity statistics from a simple ER structure can make the inference of connections more difficult. This is an important conclusion because local cortical circuits can deviate substantially from an ER structure (Song et al, 2005).

## 8 Feedforward synaptic input from unrecorded neurons can make inference more difficult

Another simplification often taken in many studies, and so far here as well, is the assumption that external input to the recorded network is uncorrelated (**Γ** and **Φ** diagonal). Local cortical circuits receive input from other cortical layers and cortical areas, and this input is likely to be correlated due to overlapping synaptic projections and due to correlations between the spiking activity in these upstream networks. Though the discriminability of these systems is no longer fully analytically tractable, we are still able to introduce some qualitative features which can impact the numerically estimated AUROC. We also introduce some additional regimes for the precision and show how sensitive the the resulting inference is to the parameters governing the non-independent external input variability.

When external input is correlated, **Φ** is no longer diagonal and we can expand Eq. (7) on an element-wise basis to obtain

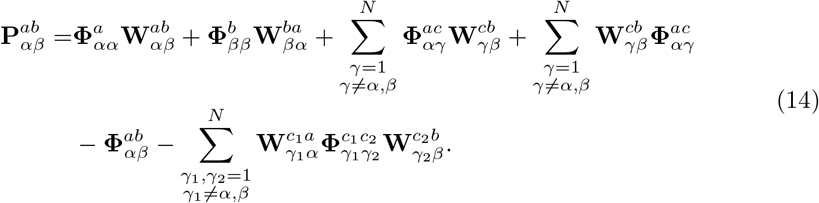

The first two terms in this sum represent precision inherited from direct connections, which are modulated by the external input precision, **Φ**. The subsequent terms represent precision inherited from common input, which is modulated by connection strength. This expansion reveals that while there remains a linear relationship between the measure and bidirectional connectivity, it is now additionally modulated by the diagonal parts of the external input covariance. The term which had previously taken the role of shared post-synaptic targets now also receives additional variability from the sources arising from shared pre-synaptic feedforward targets.

We model correlated external input as an external population of *N_x_* unrecorded neurons making random synaptic projections onto the recorded network (*i.e*., the external population is not included in the precision matrix). When correlated external input is present, we will enforce the random structure of the form

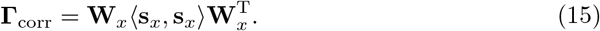

This structure models a population of *N_x_* external spike trains with the *N_x_* × *N_x_* cross-spectral matrix, 〈**s**_*x*_, **s**_*x*_〉, that sends feedforward input to the recurrent network through a random *N* × *N_x_* feedforward connection matrix 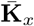, which is normalized by gains to obtain 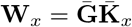 (Baker et al, 2018).

We also account for an independent noise current modeling ion channel noise and other sources of independent noise in neurons. Thus a model for the total external covariance would be

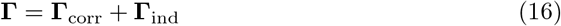

where **Γ**_ind_ is a diagonal matrix representing the variance of this additional source of independent white noise input to the system. Since **Γ**_ind_ is full rank, we may now recover invertibility of **Γ** even when *Γ*_corr_ has eigenvalues of zero, which is the case whenever *N_x_* < *N*. This model can be re-expressed by way singular value decomposition (SVD) of **Γ**_corr_ and the Woodbury matrix identity into **UDV** for some diagonal **D** to yield a linear combination of the independent and additionally modified precision values

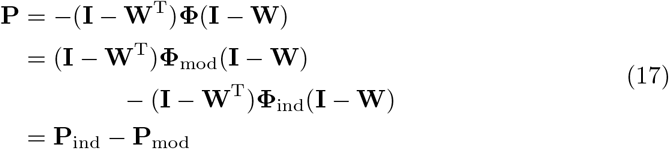

where 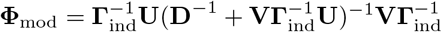 and 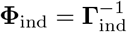. In some sense then, the information regarding network structure inherent to **P**_ind_ will remain present to some degree in the total **P** with **P**_mod_ either adding extra information about connectivity or corrupting the information from **P**_ind_ with additional noise. The first case is what is observed in the first combination case of Fig. 4 (compare red to blue). Without the regularizing independent source of external noise, discriminability is markedly decreased Fig. 4 (compare purple to others). This amplification is not ubiquitous over the parameter space, as evidenced by Fig. 4 (compare yellow to red) which causes a decrease relative to the purely independent case yet is still increased from the case of purely correlated external noise.

To understand how discriminability can be reduced by including more realistic parameters in the external network, we steadily examine each compounding source of variability beginning with the simplest. Until otherwise specified, we will begin by assuming that 〈**s**_*x*_, **s**_*x*_〉 is diagonal in Eq. (15), implying independent external spiking processes.

### Random Sparsity

Holding synaptic strengths constant and homogeneous, we will grant a random Erdos-Renyi style form of sparsity onto the feedforward projection matrix 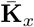. This causes additional variability since the elements of **Φ** are now random dependent on the projection structure.

### Random Feedforward Synaptic Strengths

Similar to how variability in the synaptic strengths for the recurrent layer decreased discriminability, so too may the feedforward synapses possess inherent variability in their strengths. This will necessarily induce a greater variability in the values of **Φ**.

### Correlated Spiking

We may further extend the theory to the case of correlated spiking in the external population by allowing 〈**s**_*x*_, **s**_*x*_〉 to have non-zero off-diagonal elements. This means that the external input covariance now becomes modulated by its own spiking statistics, separate from that of the randomness inherent to the network. Additionally, the scale of the external input correlations now becomes much larger: 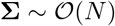 (Baker et al, 2018) compared to the uncorrelated 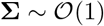 case when 〈**s**_*x*_, **s**_*x*_〉 is diagonal (Baker et al, 2018; Renart et al, 2010).

In conclusion, the relation between structure and function in the presence of latent input can depend very sensitively on both the specific model and the parameters of the unobserved network. Without any independent source of noise present in the model, the highly correlated external activity can wash out the majority of direct synaptic interactions in the recurrent network. If there is a source of independent variability for each neuron, this can help to restore and even amplify the discriminability in some cases.

**Figure 4.**
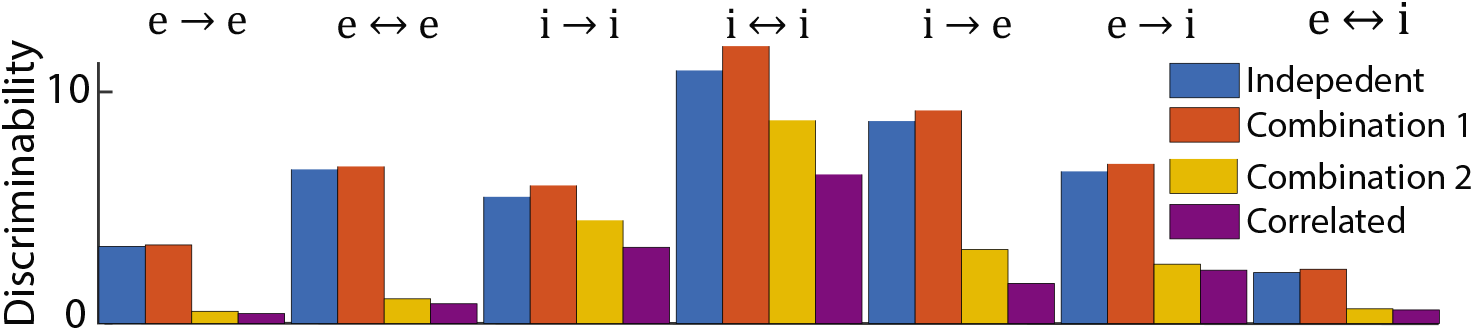
Mean discriminability across ten randomly generated networks with four feedforward connectivity types: independent (blue), a combination of rank-one and independent external input (red), a second combination of low-rank and independent input of different parameterization (yellow, see Appendix 13.5 for values), and exclusive full-rank external input corresponding to the correlated state (purple). The same recurrent networks were used across all cases.

## 9 Accounting for neuron distance or tuning differences can improve inference of connectivity

Connection probability in local cortical networks can depend on the physical distance between neurons or on their distance in tuning space, *i.e*., their tuning similarity. For data obtained by imaging methods, the lateral distance between neurons can be estimated directly. In multi-electrode array recordings, distance can be approximated by the distance between electrodes on which units were recorded (Rosenbaum et al, 2017; Smith and Kohn, 2008). Distance in tuning space can be estimated by comparing tuning curves of recorded neurons (Kohn, 2005). For example, orientation tuning difference in the primary visual cortex can be defined as the distance between neurons’ preferred orientation. These distances provide an additional type of information which can be used in conjunction with precision to improve inference of connectivity. Some intuition for this is given by an example where a distant pair of neurons is unlikely to be connected, even if their precision value is large. We next extend our theory to account for this extra source of information.

There are many variations on network models of spatial dependence. We consider a network in which each neuron is randomly assigned a preferred orientation, *θ*, and connection probability depends on the difference between neurons’ preferred orientations. Specifically,

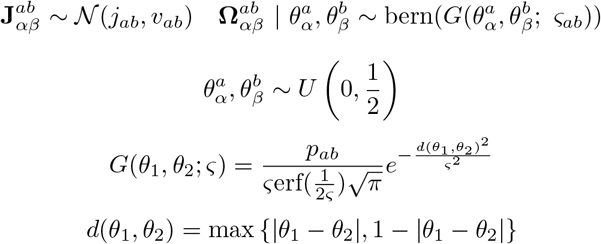

where orientations in radial units (*θ* ∈ [0, *π*] rad) have been rescaled to arbitrary units on the interval 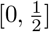 and the “wrapped” nature of the space has been maintained by way of the distance function *d*. The parameter *ς_ab_* defines the widths of the projections within or between the sub-populations *a* and *b, i.e*. the likelihood that more dissimilarly tuned neurons connect. Note that the model has unconditional sparsity levels equivalent to the standard Erdos-Renyi model. This network structure can easily be extended to two or three dimensions to model distance in physical space, yielding similar overall dependence of correlations on distance (Rosenbaum et al, 2017)

If we were to use distance alone to infer connectivity, it would give lower-quality inference for broader spatial widths (Fig. 5d), an intuitive result since very large spatial widths begin to approximate an Erdos-Renyi network in that all connections are formed with near-equal probabilities and thus the distance between pairs becomes a meaningless quantity. For certain network parameters, connectivity in each subgroup may be better inferred by one marginal measure or the other (distance or precision; Table 1), and there seems to be no simple way to decide *a priori* which metric will necessarily be better for an arbitrary choice of hyperparameters. But limiting ourselves to pairwise choice between two one-dimensional measures misses the larger implication that we are now able to classify based on the joint space of the dual measures of precision and distance seen in Fig. 5a-c,e-g,i-l. This represents our first step away from single-thresholding of measures towards the classification of connectivity by way of cluster association or linear separability in a higher-dimensional space consisting of multiple measures. This is similar to the approach used by Chambers et al (2017) to improve classification by way of an ensemble of many functional measures.

**Figure 5.**
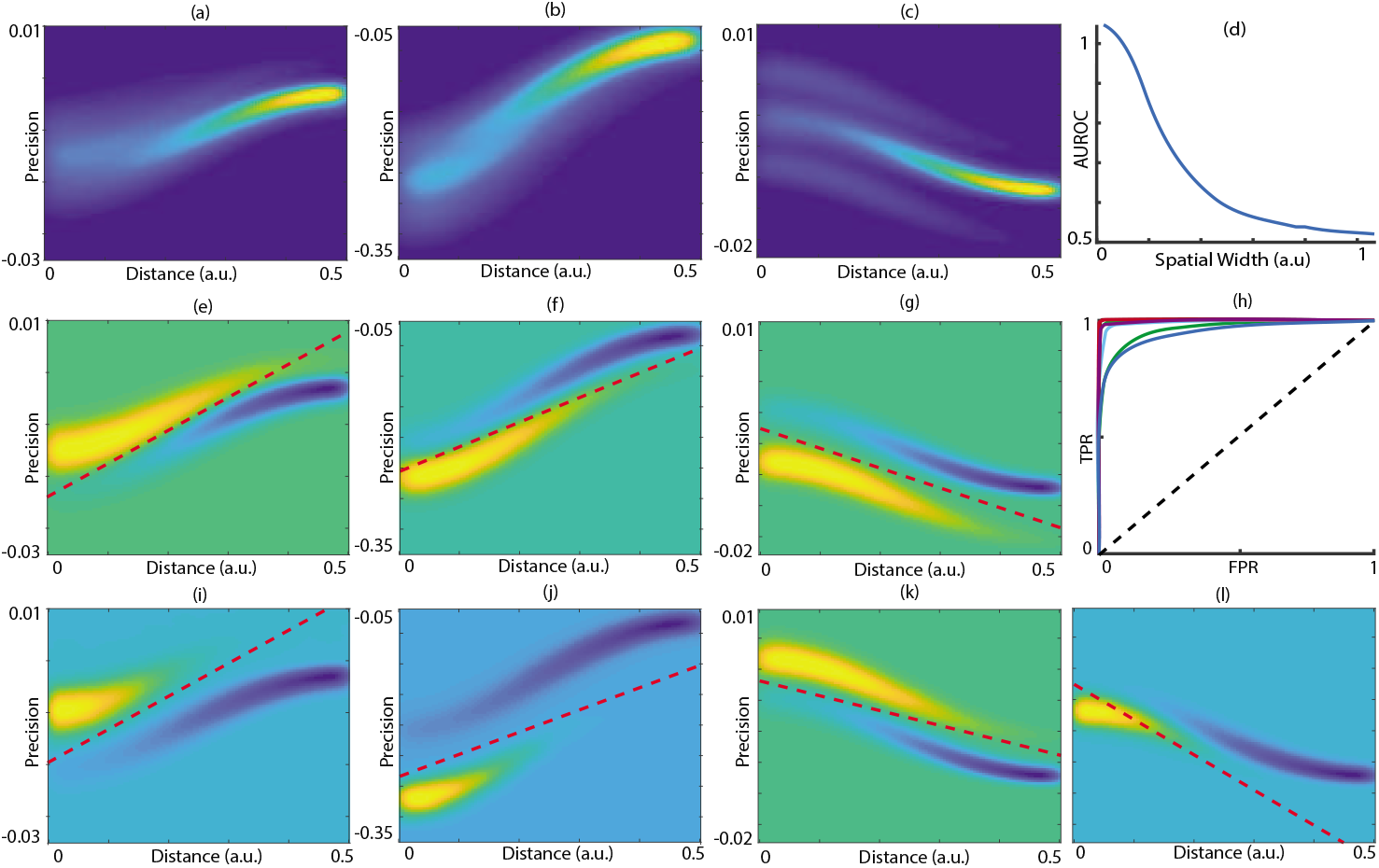
Using the knowledge of pairwise distance within spatial networks improves inference. (a-c) Two-dimensional heatmaps of the KDE of each block of a randomly generated precision structure from the spatial network type, all with unlabeled connectivity in the order: (a) excitatory, (b) inhibitory, (c) mixed. (d) AUROC for the *e* → *e* sub-population, based only on the marginal metric of distance and shown as a function of the spatial width. (e-g,i-*ℓ*) Heatmaps of the difference between the two-dimensional KDEs conditional on connection type for each subgroup assigned as follows: (e) *e* → *e*, (f) *i* → *i*, (g) *i* → *e*, (i) *e* ↔ *e*, (j) *i* ↔ *i*, (k) *e* → *i*, (*ℓ*) *e* ↔ *i*. Dashed red lines denote a linear classifier corresponding to the ROC curve in (h), with threshold fixed at the point where the sum of the number of true and false positives (*i.e*., assigned connections) equals the total number of condition positive (*i.e*., actual number of connections in the network). (h) Optimal ROC curves for each subgroup over the joint space. AUROCs in (h) are reported in Table 1.

**Table 1.**
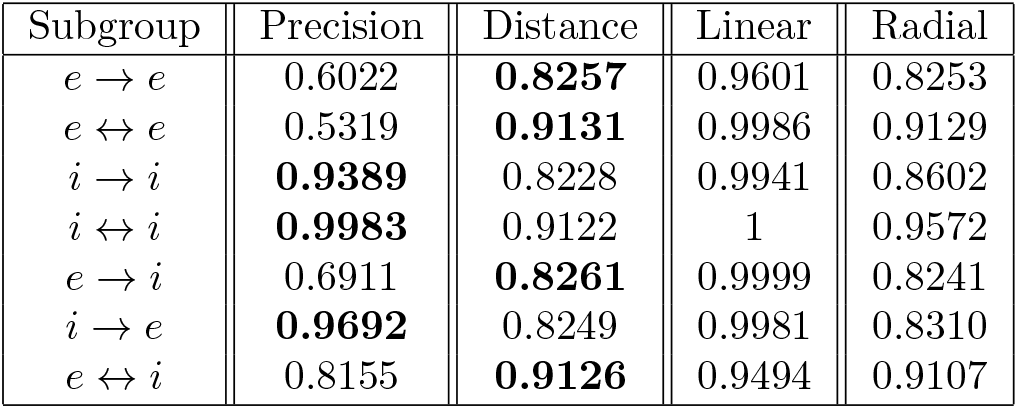
AUROC for each subgroup using each measure only in the marginal sense. The greater of the two values is emboldened for visibility. The “Linear” column of values are the AUROC of the curves seen in Fig. 5h. The “Radial” column of values are the AUROC of the curves seen in Fig. 6h.

In analyzing the use of precision and distance as dual measures for classification, we will examine the parameterized family of ROC curves induced by classifiers based on a linear combination of the two measures. That is, we define a threshold method analogous to (Maswadeh and Snyder, 2012)

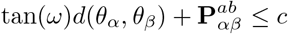

to classify connectivity, where *c* spans the entire classification space for each fixed angle *ω* ∈ [0, *π*], thereby generating ROC curves parameterized in terms of this angle. This inequality can be interpreted as follows: Draw a line in the joint space of distance and precision with slope tan(*ω*) and intercept *c* (see dashed lines in Fig. 5). Pairs of neurons with precision-distance values above this line are classified as connected. It is useful to examine the dimensional reduction of the family of curves by way of the AUROC for each slice as a function of the slope of the partitioning line; these curves are displayed in Fig. 6a-g. They primarily reveal how, for certain subgroups, the “optimal” classifier (*i.e*., the peak of each curve) may be dangerously close to the worst linear classifier possible over the space (*i.e*., the minimum of the curve). Some groups may also plateau in a more stable fashion than others, implying robustness across sub-optimal angles.

**Figure 6.**
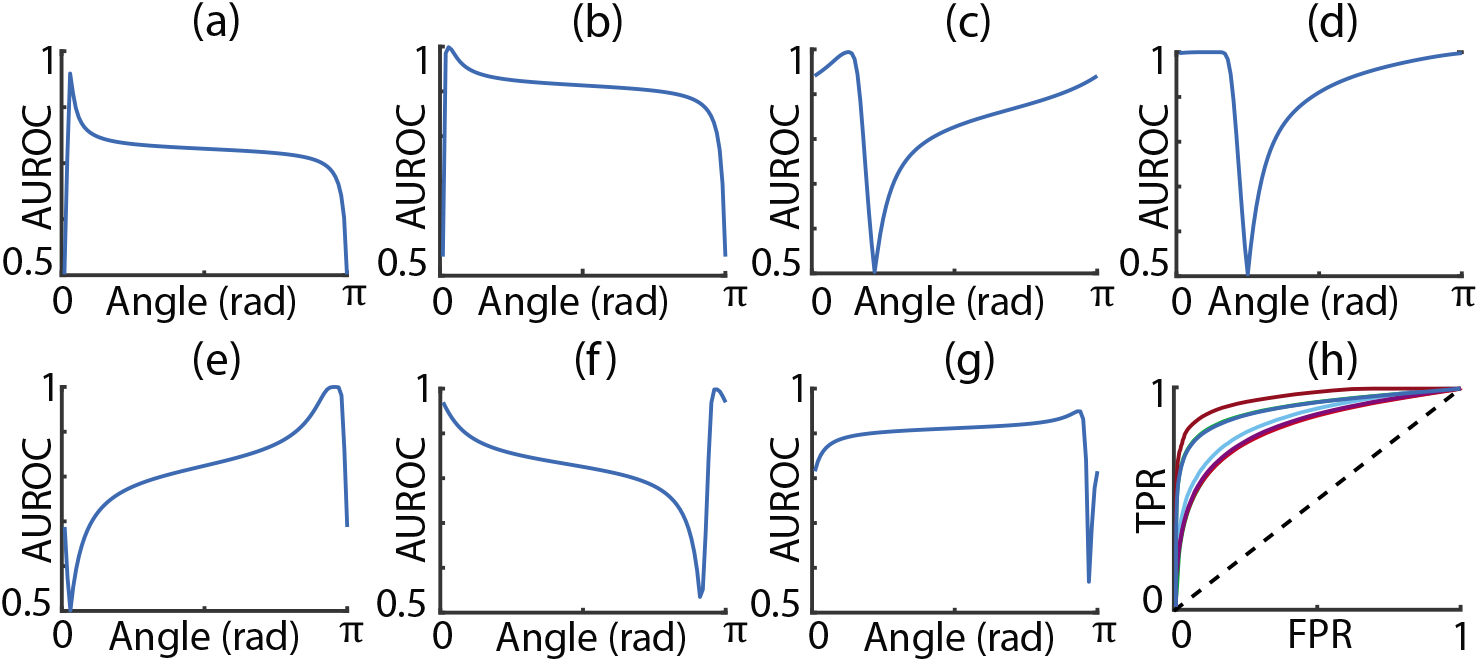
AUROC varies with respect to the angle of the projections. (a-g) AUROC as a function of the slope of a linear classifier over the joint space of precision and distance from Fig. 5. (h) ROC curves for the radial heuristic. AUROCs for (h) are reported in Table 1.

It should be noted that the marginal metrics represent the perfectly vertical and horizontal slices of this space and thus the family of curves further generalizes all higher-dimensional linear classification methods, which can include some unsupervised clustering methods such as the *k*-means algorithm. Most notable from this approach is that there exist several cases where the conditional marginal distributions of neither precision nor distance are themselves perfectly separable and yet their clusters in the joint space are almost perfectly linearly separable.

While this result is encouraging towards the use of the joint space for classification, it should be mentioned that common unsupervised methods do not work very well due to the non-linear and non-Guassian relationship between precision and distance. Whilst supervised methods can easily learn this relationship, these lack applicability to *in vivo* data where ground truth, and indeed the true joint relationship, remain unknown.

To alleviate the difficulty in choosing either a marginal measure or the slope for a linear classifier, we further introduce a basic heuristic which would be immediately applicable to real data in which both precision and distance are inferable. Our heuristic classifies connectivity (or anti-connectivity) as the points which lie on the interior of a circle centered at the peak of the unlabeled two-dimensional kernel density estimate (KDE). More precisely, a pair of neurons would be classified as connected if

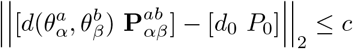

where || · ||_2_ is the Euclidean norm, and [*d*_0_, *P*_0_] is the location of the peak (the argmax) of the empirical KDE. Hence a pair of neurons is classified as connected if their precision-distance value lies inside a circle of radius *c* centered at the peak of the estimated joint density of distance-precision values. This method is also mathematically equivalent to directly thresholding the likelihood of points transformed into a Gaussian/radial basis with an identity covariance. But unlike the angular method, this produces a single ROC curve without relying on the choice of any other parameters (like *ω* from above), though further improvement would undoubtedly be gained from specifying or optimizing the covariance relation between parameters within the Gaussian basis.

We assess this heuristic by varying the radius, *c*, of the circle to span the classification space, generating the ROC curves in Figure 6h. As observed in Table 1, the AUROC obtained from this heuristic is always less than the best marginal measure, but always much better than the worst one. It also tends to mimic distance as a measure in this regard, being very similar in quality when distance is the preferred measure. Thus the heuristic is convenient in that it removes blind choice between marginal measures and the arbitrary choice of a parameter like *ω* above, even though it may sometimes be suboptimal.

Our conclusion for spatial networks is encouraging - despite the increase in precision variability due to the spatial configuration, the joint use with known distances greatly improves the overall quality of inference and has direct application to *in vivo* recordings.

## 10 Inferring connectivity from network simulations

Up to this point in our study, we have only quantified the quality of inference under an assumption that we have a perfect estimate of the precision matrix. This will not be the case for actual data generated from explicit dynamics, be it *in silico* or *in vivo*. An account of the additional variability from imperfect statistical estimation using inversion of the covariance matrix is now given for Gaussian data reflecting the structure of Eq. (8) using Erdos-Renyi networks with independent external input. Our analysis will show that in order to achieve near-optimal levels of inference (near-optimal being relative to what the system itself constrains the maximum theoretical AUROC to), very large sample sizes are required, corresponding to experimental recording lengths that may not always be feasible in practice.

This analysis yields a total discriminability of

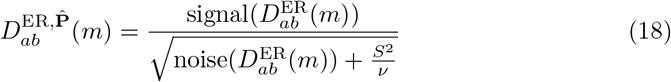

where *ν* is the number of samples used to estimate the precision matrix, which is proportional to the duration of a recording. The *S*^2^ term relates to the additional variability from the statistical sampling, with an explicit form found in Supplemental Section 6. As expected, this new form indicates that inference on the estimate is always less than the perfect case but increases monotonically with the number of data points governed by the length of a recording or simulation.

ER P

Note that as the number of samples, *ν*, tends to ∞, 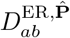 tends to the “optimal” discriminability 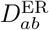 derived under the assumption of a perfect estimate of **P**. This is the value of discriminability discussed in previous sections and represents the most one can recover about connectivity from precision, but does not necessarily represent perfect recovery of connectivity.

An interesting perspective is offered by solving the full discriminability equations to obtain a direct relation between the number of samples *ν*_0_ required to obtain a target fraction, *ϕ*_0_, of optimal discriminability

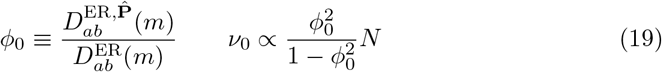

which illustrates two important concepts: It grows linearly with the number of neurons (*N*) and the nature of the rational function in *ϕ*_0_ necessitates disproportionately larger values of *ν*_0_ (more samples) to achieve higher optimal percentages.

This theory is explored first using simulations of an OU process like the one defined in Eq. (1) whose parameters are the same as the top row of Fig. 1. Estimates of inverse sample covariance were obtained at regular intervals and used to compute the convergence rates in Fig. 7a. The final ROCs obtained are then displayed in Fig. 7b. Since the network used in the OU simulations was the same as in the majority of Fig. 2, the same nearly ideal ROC curves should be observed as sample size tends to ∞. Thus all suboptimal ROCs observed in Fig. 7b are the result of finite sampling of the precision matrix. Further, the rate of convergence in Fig. 7a is consistent with the rational expression in Eq. (19). The rate of convergence for *ϕ*_0_ is also radically different for each subgroup in Fig. 7a, a feature also explained analytically by finding the constant of proportionality in Eq. (19) as a function of the sampling variability *S*^2^, derived in Supplemental Section 6. Of particular importance then is the bulk of connectivity contained within the excitatory population, which due to its relatively weak synaptic proportions (see Appendix 13.5) produces very small correlations, which in turn requires many more samples to adequately estimate.

**Figure 7.**
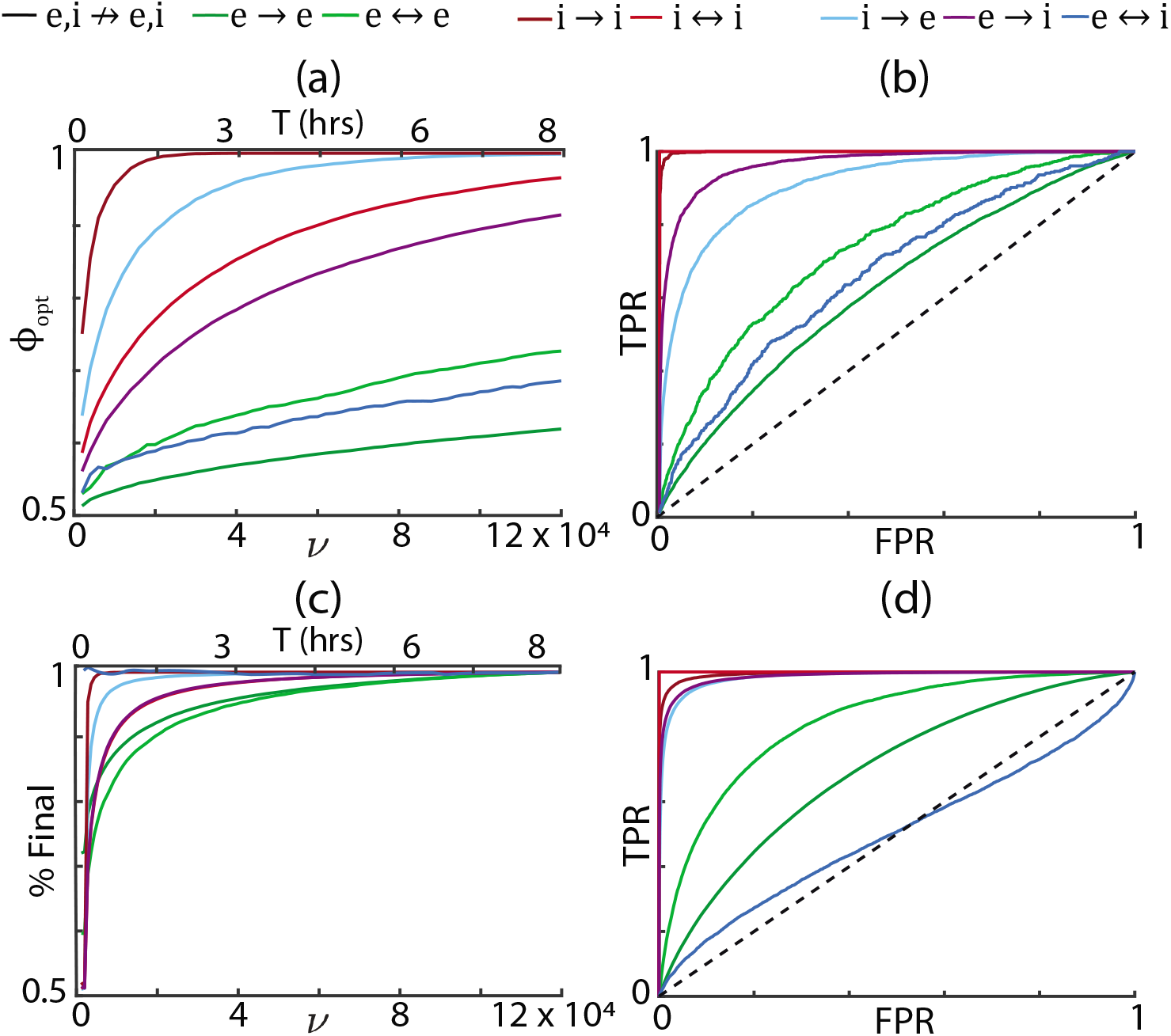
The rate of convergence to the maximum obtainable AUROC depends on neuron model. (a) Plot of mean *ϕ*_0_ versus number of samples (*ν*) for four networks of *N* = 500 neurons following an OU rate process with independent noise. (b) The ROC curves of the final point in time of the simulation. (c) Plot of the AUROC as a function of time, normalized by the final endpoint, for a network of AdEx spiking neurons in the correlated balance regime. (d) ROC curves of the final point in time of the simulation. In both (a) and (c), the top axis of equivalent time in rounded hours is shown for comparison to *ν*, using an integrating time window of 250 ms. It should be noted that the timescales for the OU process in (a) are subjective and may not map directly to biophysical recordings; whereas the timescales of the EIF in (c) are chosen to be realistic and time may be interpreted directly.

Yet another source of variability arises if, instead of assuming a linear Gaussian model (OU process), we consider a non-linear model such as that induced by a large balanced network of adaptive exponential integrate-and-fire (AdEx) neurons, defined by the membrane potential dynamics

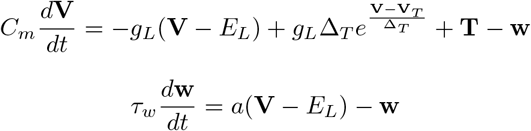

subject to the rule that if a voltage exceeds the threshold 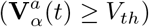 then it returns to reset 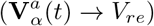, its adaptation current is incremented 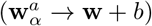, and a spike is recorded. The input terms stemming from recurrent sources **R** and external sources **X** may be expressed as

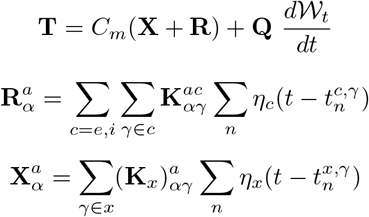

where 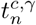 is the *n*^th^ spike time of neuron *γ* in population *c* = {*e*, *i*, *x*} and synaptic kinetics are modeled by the filter 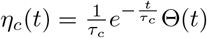 where Θ(*t*) is the Heaviside step function. This network is held in the aforementioned correlated state by letting the external spike times *t^x^* be correlated across pairs of feedforward projecting neurons, giving 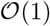 mean spike train correlations in the recurrent network (Baker et al, 2018). This scaling allows spike count covariance to become much stronger than those produced in the case of OU processes (Baker et al, 2018), leading to more accurate estimation of precision and therefore better recovery of connectivity (7c).

As we have now transitioned to a spiking model of neuron activity, we must adjust our notion of covariance to be taken over spike counts aggregated over time windows of moderate size (~ 250 ms). By aggregated spikes over time windows larger than the decay of their autocorrelation, we begin to approximate the zero-frequency structure of Eq. (6) and the results implied by its inverse through the previous sections.

However, the optimal AUROC in 7d is still hampered by at least two new sources: i) the non-linear characteristics of the model impart a certain deviation from the approximating equations, and ii) even within the linear approximation, the gains for each neuron (derivative of *f*-I curve) are no longer fixed parameters – they become random variables following a distribution with non-zero variance. While the variability coming from the gains reduces inference, the mean value may actually act to improve it beyond the simpler cases previously considered. To exemplify, note the parameters of the balanced network use identical synaptic proportions to those in the OU process, but the magnitude of the average synaptic strengths is nearly a hundred times stronger when measured in consistent units. In the previous qualitative analysis this would imply near-zero discriminability, yet that considered only gain values fixed at unity. In the AdEx simulation, the gains were estimated empirically by a quadratic function fit to the *f*-I curve, similar to (Ebsch and Rosenbaum, 2018). When compared in consistent units, the corresponding synaptic strengths were found to be small on such an order than returns the product 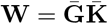 back to a level we would expect reasonable discriminability. In short, stronger recurrent synapses become modulated by weaker gains, enabling a wider range of network parameters to be viable in conveying structure via function as measured by precision of spike count covariance over large time windows.

In practice, we recognize that precision matrices are more often estimated by way of far more rigorous regularization techniques such as graphical-LASSO, shrinkage, or sparse-latent (Yatsenko et al, 2015) methods to improve the quality of estimation for small quantities of data by utilizing the assumed sparsity of the matrix. Unfortunately, these methods do not allow simple analytical properties such as the variance of the estimator to be inferred and so we use the general estimator here to establish an upper bound for the sample size *ν* or, equivalently, the recording length of T hours using the scale proportion *T* ≈ 7*ν* × 10^-5^ using integrating time windows of 250 ms. It is then assumed that proper use of regularization (*i.e*., ideal choice of regularization strength) would offer a reduction in this variance equating to smaller bounds on sample size necessary to achieve target levels of inference.

In conclusion, accurate statistical estimation of precision and the ensuing use of the measure for inference of structural connectivity remains a very hard problem. If a given neural region exhibits strong correlations driven by external variability then it may be possible to reach asymptotic levels of inference with relatively short recording times, but the nonlinearities of that regime diverge from the analytical theory developed in previous sections and may lead to suboptimal inference within certain sub-populations.

## 11 Mean-Field Analysis of *In Vivo* Data Suggests Exclusive Sources of Input for Inhibitory Sub-Populations

Following the analysis of inferring pairwise connectivity from the precision matrix, we are left with the question of what exactly can be inferred using *in vivo* data sampled at low temporal resolution and at recording durations too short for accurate estimation of the entire precision matrix. One option left to us is to examine what can be inferred from the mean-field statistics of the neuron cell type sub-populations, *i.e*., from the cell-averaged values of each block in **Σ**. These results will provide some interesting insight into the mean-field structure of external projections onto an observed recurrent layer.

A mean-field theory of correlated variability in balanced networks shows that for large *N* (Baker et al, 2018; Renart et al, 2010; Rosenbaum et al, 2017)

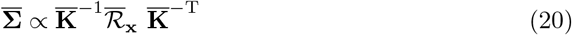

where

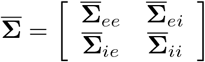

is the 2 × 2 matrix of cell-type averaged covariances with

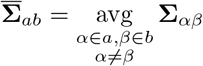

and similarly for the 2 × 2 mean-field external input covariance matrix, 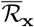. The 2 × 2 mean-field connectivity matrix is defined similarly with

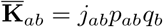

where *q_b_* = *N_b_*/*N* is the proportion of neurons in population *b* = *e*, *i*, and we remind that *j_ab_* is the mean synaptic strength of projections from *b* to *a* which occur at a mean connection probability of *p_ab_*. Importantly Eq. (20) is independent of the gains that were present at the pairwise level and implies then that

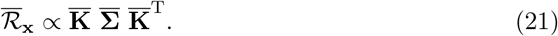

When a large number of cells are recorded from a short-duration recording, 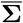 can be estimated more accurately than the full matrix **Σ**. The mean-field connectivity, 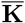, still cannot be inferred without knowledge of 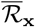, which is typically not known in experiments. However, the connection probabilities, connection strengths, and sub-population ratios that define 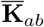 have been estimated from intracellular recordings (Jiang et al, 2016). This allows us to solve a reversed problem: Instead of inferring mean connections, 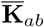, we can combine estimates of 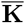 from intracellular recordings with estimates of 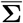 from multi-cellular recordings in the same cortical area to obtain an approximate estimate of external input covariance, 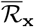, by directly applying Eq. (21).

Specifically, we set *q_e_* = 0.8 and *q_i_* = 0.2 then used measurements of the maximal evoked post-synaptic potential (units mV) from intracellular recordings of connected pairs of Pyramidal and Basket Cells in L2/3 of mouse primary visual cortex (Jiang et al, 2016) to constrain

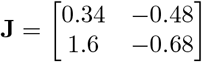

together with their corresponding estimates of connection probabilities from (Jiang et al, 2016)

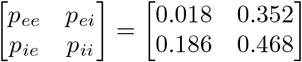

Note that for any model with a single homogeneous population of neurons providing external synaptic input to the recurrent network, (including the previous AdEx spiking network), the 2 × 2 matrix, 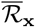, of population-averaged external input covariances is proportional to

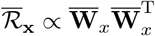

where 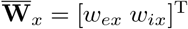 is the 2 × 1 mean-field feedforward connectivity matrix defined similarly to 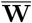 (Baker et al, 2018; Renart et al, 2010; Rosenbaum et al, 2017). As a result, 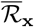 is rank one (and therefore has determinant zero) for any such model. Hence, the product of the off-diagonal elements of 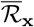 should be equal to the product of its diagonal elements. We may therefore test the hypothesis that the recorded network receives correlated external input from a single homogeneous population of neurons by comparing the product of the off-diagonal to the product of the diagonal elements of the estimated matrix 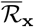.

We proceed to analyze a dataset of 11 recordings on 5 individual mice. In each recording session, between 163-385 neurons were recorded via 2-photon calcium imaging of mouse primary V1 cortex L2/3. Each recording consisted of around 200 trials per presentation of 2 stimuli consisting of lines oriented at either 0° or 90° angles. The fluorescence traces from each trial were then deconvolved using the fast non-negative deconvolution of Vogelstein et al (2012). For more details on experimental methods, please consult Appendix 13.1. The covariance between neuron pairs at each point in time was then calculated across trials for each stimulus type, and subsequently averaged over time in order to extract the noise, rather than stimulus, covariance. Non-firing neurons were dropped from the covariance estimation step. The mean-field was then taken over each covariance matrix in each stimulus/experiment, using the parvalbumin (PV) labeling to define the excitatory (PV−) and inhibitory (PV+) cell types. We note that not all PV− types are necessarily excitatory, as there are many types of inhibitory interneurons in this region which are PV−, but the sampling probability of these should be low enough to not significantly affect our analysis.

These mean-field averages of the resulting noise covariance matrices for each experiment are seen in Fig. 8c and are subsequently passed through Eq. (21) using the previously specified values to constrain the degrees of freedom. This results in the values shown in Fig. 8d which do not appear to obey a rank one property over all subjects. Treating each point in Fig. 8e as a sample, we perform a simple hypothesis of mean-equivalence using the Behrens-Fisher test to verify this result. The null hypothesis that

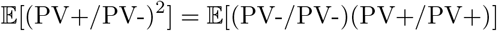

is rejected with a *p*-value of 0.0143 and thus we cannot claim that the mean-field external input covariance for mouse V1 cortex L2/3 is rank-one. There are multiple candidate models to explain the additional rank but one of the simplest is the existence of exclusive input to the PV+ sub-population illustrated in Fig. 8b, and first proposed by (Yatsenko et al, 2016).

**Figure 8.**
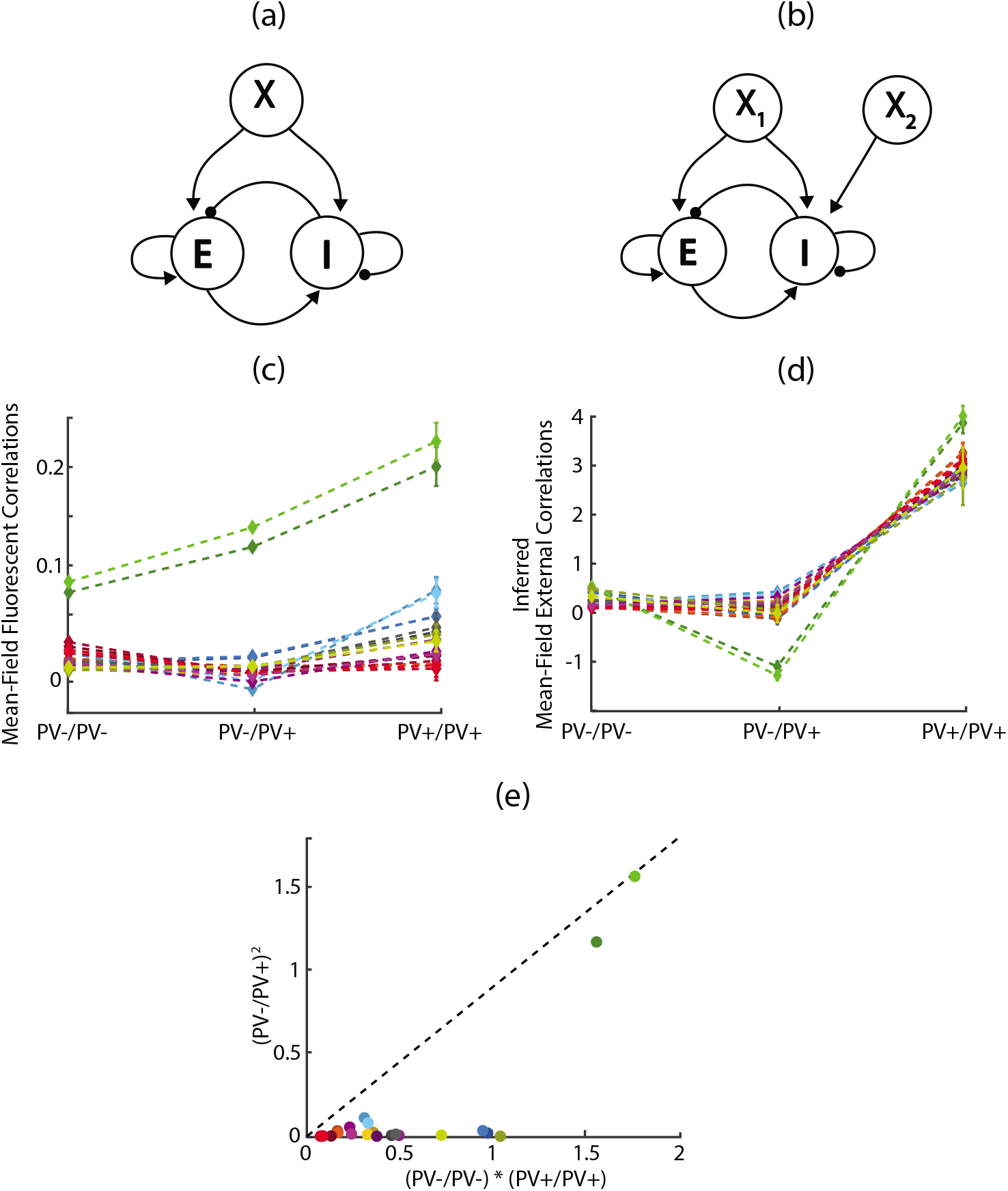
In vivo recordings indicate a deviation from an assumed model of the external input. (a) Originally hypothesized model of external input to the recurrent network, consisting of a single homogeneous population. This is the structure used in previous EIF simulations. (b) Alternate model to explain observed high-rank structure of *in vivo* data, which uses two distinct external populations with one projecting exclusively to inhibitory cell types. (c) Estimates of mean-field noise correlations partitioned by parvalbumin containing (PV+) and non-containing (PV−) subtypes over 12 total experiments (each a different color), with standard error bars shown around each experimental mean-field (but only visible for the PV+/PV+ blocks). (d) Estimates of mean-field external input covariance using values in (c) in conjunction with Eq. (21) and intracellular estimates of synaptic values. (e) Plot of the product (PV−/PV−)(PV+/PV+) versus (PV+/PV−)^2^, where each quantity is scaled by the arithmetic mean of the three values to maintain visual perspective. Rank one matrices would have all values close to the diagonal reference line.

There may be a question of how sensitive our conclusions are if we consider the variability in estimating the mean values for *j_ab_* and *p_ab_*. Thankfully, Jiang et al (2016) report standard deviations for the post-synaptic potentials as well as the number of neuron pairs recorded for the connection probabilities. Both of these can then be used to construct a Monte Carlo approximation to the total contributed variability in the final estimates of 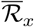. We find that all perturbed versions of Figs. 8c-e do not qualitatively change in any significant way, and indeed the distribution of p-values for the previous statistical test is largely within a range of less than 0.1 (Supplemental Figures 2–5).

## 12 Discussion

It remains an open question as to how our theory interacts with subsampling (Brinkman et al, 2017; Paninski, 2004; Pillow et al, 2008), where external correlations are caused not by correlated external spike trains but instead by latent (unobserved) recurrent interactions. From an analytical perspective, the term **Γ** which occurs in the inner part of Eq. (6) would take a different form to incorporate network-valued functions relating the observed and unobserved parts, rather than simply being a parameter of the system. While intuition indicates that recovery performance should scale increasingly with the proportion of the recurrent network observed, this has not been shown using our functional measure nor our biophysical models. This would be important to a future application of this research to *in vivo* results, as it is uncommon to have techniques capable of recording from an entire self-contained recurrent network.

There are many ways in which the theory we have established could be improved by statistical methods. For example, in estimating the sample precision we directly inverted the covariance matrix, which is not a commonly used method, but we chose it to glean analytical results for the unconditional variance of the model. While this ought to serve as a lower bound for more accurate methods such as the graphical-LASSO family or neighborhood based methods, it is unknown if there is an upper bound on how well such numerical estimations could improve the quality of inference.

A main implication of our results is that knowledge of cell-types is extremely useful in untangling the full mixture distribution. In this paper, we only distinguished primary excitatory and inhibitory cell types in the two-population model; real neural circuits contain a variety of neuron types and subtypes with intricate connectivity properties (Jiang et al, 2016; Pfeffer et al, 2013). A more realistic model of a real system then would be a multi-population network structure.

We performed much of our analysis in the limit of large network size assuming a 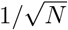 scaling of synaptic weights. Asymptotically larger synaptic weights typically violate stability conditions on the network dynamics at large *N*. Some studies have considered sparsely or weakly coupled networks (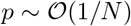 or 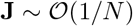). Our analysis can be modified to these settings. In principle, inference of synaptic connectivity from precision becomes perfect in the limit of large network size under these scalings when one has a perfect estimate of the precision matrix. However, asymptotically weak coupling in this case implies that asymptotically longer samples (as in Eq. 19) are required for accurate estimation of the precision matrix in this limit, suggesting that inferring synaptic connectivity in weakly coupled networks is difficult in practice.

Even aside from functional-effective measures and ensembles thereof, there is also modern work showing that introducing large targeted perturbations to nodes within the network, and assigning connectivity based on observed responses throughout system (Widloski et al, 2018). While this has been shown to give good recovery for some *in silico* regimes, it remains unknown how it depends on network parameters similar to what we have examined. Another aspect of these perturbational methods is the experimental difficulty that would be required in application. While it is possible to use intracellular stimulation in conjunction with both calcium imaging and micro-electrode arrays, the scalability of the perturbational methods would be impractical for networks on the order of thousands of neurons.

Another side topic of this paper is in regards to sub-optimal thresholding methods in ROC generation. Specifically, whenever the ROC curve dips below the diagonal, there is indicated loss of information due to a non-monotonic likelihood ratio between the two compared distributions. The ideal method would then be to correct the thresholding based on knowledge of this relationship, seen in Supplementary Figure 1. Without this knowledge, a simple way to enforce monotonicity is to take the absolute value of the points being thresholded, much as we did in Fig. 1h. Note however that doing so makes the underlying mixture distributions non-Gaussian and so discriminability analysis only serves as a lower bound on performance. Use of such transformations in real data would ultimately be a subjective choice, though possibly informed by in silico results similar to this paper; nonetheless, it is difficult to justify without making prior assumptions on the structure of the real data.

In addition to the radial heuristic we proposed for inferring connectivity from precision and distance measurements, the ROC curves of which are seen in Fig. 6h, there are undoubtedly many other heuristics may consider either alternate centerings or more complicated geometries such as ellipses to account for the correlation within the measures and these could certainly do better than ours. But allowing more degrees of freedom to the model also increases its subjectivity; we illustrate ours simply for the sake of example to show how the transition into the higher-dimensional data space may allow for more detailed thresholding heuristics than exist in the univariate case.

Finally, some authors who perform *in silico* benchmarking refrain from use of ROC curves and instead favor precision recall (PR) curves, accuracy curves (ACC), or their own custom metrics. It should be noted that PR curves may only be reliably used under very sparse settings, and even then may only compare the relative performance of different methods on an identical network. Notably, it is not valid to vary parameters implicit to the model and examine how a metric such as the area under the PR curve changes as a result. Likewise, ACC curves are imbalanced to unequal class sizes, and will show misleading recovery results in the presence of high true negative to false positive ratio.

## 13 Appendix

### 13.1 Experimental Methods

All procedures were carried out in accordance with the ethical guidelines of the National Institutes of Health and were approved by the Institutional Animal Care and Use Committee (IACUC) of Baylor College of Medicine.

The animals (*n* = 5 PV-Cre/Ai9 crosses on a C57Bl/6 background, labeled with the fluorescent marker tdTomato) with an age range of p40 to p60 were initially anesthetized with Isoflurane (3%) and then anesthesia was maintained by either Isoflurane (2%) or a mixture of Fentanyl (0.05 mg/kg), Midazolam (5 mg/kg), and Medetomidin (0.5 mg/kg) with anesthesia boosts consisting half of the initial dose every three hours. The body temperature of the animal was maintained at 37C throughout the surgery using a homeothermic blanket system (Harvard Instruments). In some experiments we applied eye oil ointment (polydimethylsiloxane) to prevent dehydration of the cornea. Surgery and dye injections of the Oregon Green 488 BAPTA-1 AM (OGB1, Invitrogen) calcium indicator were performed as previously described (Garaschuk et al, 2006).

We used stereotactic information to locate our recordings to the primary visual cortex of the mouse (V1). In some experiments we used intrinsic imaging to verify the location of V1 (Kalatsky and Stryker, 2003). We recorded calcium traces using a custom built two-photon microscope equipped with a Chameleon Ti-Sapphire laser (Coherent) tuned at 800 nm and a 20x, 1.0 NA Olympus objective. Scanning was controlled by a custom built acousto-optic deflector system (AODs) (Cotton et al. 2013). The average power out of the objective was kept less than 120mW. Calcium activity was typically sampled at a mean of 260Hz (min/max: 78-450 Hz). We recorded data from depths of 100-540 *μ*m below the cortical surface.

The measured fluorescent traces were preprocessed in order to reduce common mode noise related to small cardiovascular movements (Cotton et al, 2013) and the firing rates were estimated using by nonnegative deconvolution (Vogelstein et al, 2012).

### 13.2 OU Identity for Normal 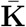

For the OU system defined in Eq. (1), if **KK**^T^ = **K**^T^**K, G** = *g***I**, and **QQ**^T^ = *σ***I** we have

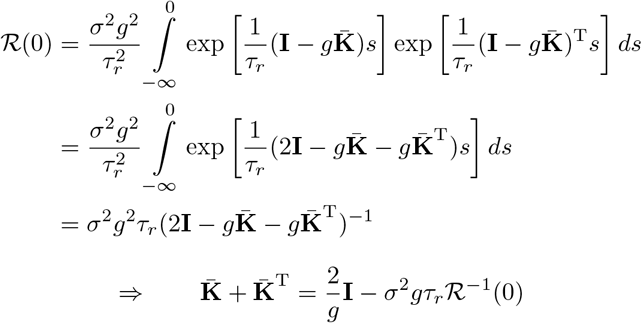

### 13.3 Relation Between AUROC and Discriminability for Normal Distributions

It is well known that the area under an ROC curve may be parameterized and solved to yield the identity

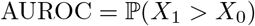

for normally distributed scores in the positive and negative classes *X*_1_ and *X*_0_, respectively. If these scores are normally distributed then closure properties imply

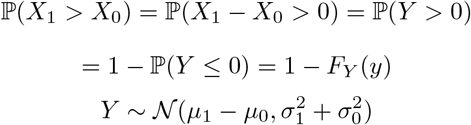

and so

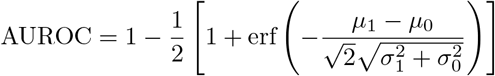

and so by appropriately defining the discriminability *D* and re-arranging we obtain

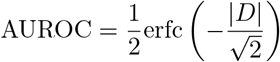

where the absolute value restricts the AUROC to 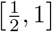 which corrects for anti-classifiers. The result is the bijective mapping from *D* to AUROC, leading to the natural invertibility between the two which is necessary for our arguments.

### 13.4 Central Limit of Precision with Independent External Input

Precision values from the case of independent external input can be expressed element-wise as

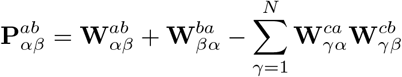

which in the thermodynamic limit of network size (*N* → ∞) leads the summation term to converge to the limiting normal distribution, so long as all elements of **W** are independent with finite variance. So long as we enforce normal distributions on and zero-variance gains on the synaptic strengths, and since the normal distribution is closed under summation, elements of **P** will also be normally distributed. All these assumptions hold under the standard ER case as well as the CER case, whereby correlations are specially constructed so that pairs of the form 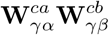 are independent across *γ* though not across pairs of *α, β*. The only breakdown of the CLT independence conditions occurs in the HD_out_ case, as all such pairs are now highly correlated across *γ*, giving an apparent power law limiting distribution instead.

While the above requirements on *J* hold exactly only if it is normally distributed, as long as **Ω** is Erdos-Renyi this theory will still hold as a fair approximation since the 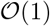 part of the precision distributions are determined by the summation term with the direct strengths offering only a 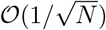 deviation. Even if synaptic weights are specified from a one-sided distribution of finite variance, the induced asymmetries against the limiting Gaussian will dissipate in the large *N* limit. Any variance present in the gains will also lead to small errors with an exact Gaussian, but these effects will again decay for large *N* and so the discriminability theory outlined in the paper should still hold as a good approximation for the expected AUROC.

### 13.5 Simulation and Figure Parameters

For all simulations, we use an alternative parameterization of synaptic strength which makes modulation easier to control. We express the average synaptic weights as

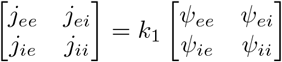

where the *synaptic proportions ψ* are normalized by the inhibitory component as *ψ_ab_ = j_ab_/j_ii_* and the mean *synaptic magnitude k*_1_ is then modulated while holding the proportions fixed. Similarly, the synaptic variance is parameterized as

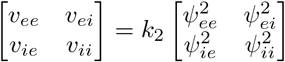

where the magnitude of *synaptic variance k*_2_ is modulated. For all figures and simulations, we use a version of the synaptic proportions used in Pyle and Rosenbaum (2016) that have been perturbed in order to give non-zero real part to the eigenvalues. These proportions are

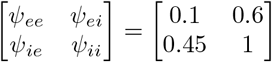

as well as the fixed synaptic ratios of *q_e_* = 0.8, *q_i_* = 0.2 and the same fixed density of *p_ab_ = p* = 1.

In Fig. 1, we used *σ_a_* = *σ* =1, *g_a_* = *g* =1, and *u_a_* = *u* = 0 for both rows in order to simplify the interpretation. For the top row we took *k*_1_ = 2.5, *k*_2_ = 6.25 × 10^−5^ and for the bottom row, *k*_1_ = 12.5 and *k*_2_ = 1.25. The same exact networks used in Fig. 1 are used in Fig. 2, though they are partitioned according to the more informative mask.

In Fig. 3, all shared network parameters are identical to Fig. 1, except *k*_1_ = 2.5, *k*_2_ = 1.25. The model specific parameters are *ρ* = 0.2 for the CER model and *μ* = 5, *ξ* = 0.25 for the HD models. As mentioned, *σ* for the HD model is estimated numerically using the bisection method to find the fixed point.

In Fig. 4, the same recurrent networks were used across all cases. Shared recurrent parameters are consistent with the top row of Fig. 1. Shared external input parameters are *p_ax_* = *p_x_* = 0.1, *j_ax_* = *ψ_ax_k*_*x*,1_, 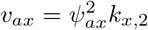, *ψ_ex_* = 1.333, *ψ_ix_* = 1. Individually modified parameters are *k*_*x*,1_ = 0.2571, *k*_*x*,2_ = 0.6571, 〈*s_x_*, *s_x_*〉*_αα_* = *r_x_* = 10, 〈*s_x_*, *s_x_*〉*_αβ_* = *c* = 0.1 for *α* = *β* in both the full-rank correlated state and the correlated part of the first combination. The first combination also had **Γ**_ind_ = **I** for the independent part. The second combination used *k*_*x*,1_ = 250, *k*_*x*,2_ = 0, *r_x_* = 10, *c* = 0, *q_x_* = 0.2, **Γ**_ind_ = 10**I**. Full-rank effects in the correlated case are induced by taking *q_X_* = 7 to give *N_X_* = 14,000 total external neurons, making **Γ** invertible with high probability even without regularization from the independent external source.

In Fig. 5, shared parameters are consistent with the top row of Fig. 1 and used a spatial width of *ς_ab_* = *ς* = 0.2. The additional spatial variability inherent to these networks accounts for the difference in AUROC using only precision as a marginal metric between Fig. 2 and Table 1.

In Fig. 7, network parameters for the OU model are identical to the top row of Fig. 1 and have independent noise level *σ* = 0.1 and timescale *τ_r_* = 1, discretized at the level of *dt* = 0.1. Shared network parameters for the EIF model are the same except for *k*_1_ = 250 and *k*_2_ = 0, as well as the fact that the statistics of the gains are no longer specifiable and are a consequence of the non-linearity of the system. AdEx-specific parameters are as follows: *C_m_* = 1, *g_L_* = 0.0667, *E_L_* = −72, *V*_th_ = −50, *V*_re_ = −75, Δ*_T_* = 1, *V_T_* = −55, *τ_w_* = 150, *τ_e_* = 8, *τ_i_* = 4, *τ_x_* = 10, *a* = 0, and *b* = 0.1. External input followed: *q_x_* = 0.2, *k*_*x*,1_ = 250, *k*_*x*,2_ = 0, and identical *r_x_*, *c*, *ψ*, *p_x_* as from Fig. 4 purple. The balanced network exhibited less than 20% relative error to both the balanced rate approximation and the mean-field covariance approximation from Baker et al (2018).

## Acknowledgments

We would like to thank Ryan Pyle (Notre Dame) and Krešimir Josić (University of Houston) for feedback and many productive discussions on this topic.

## Supplemental Material

### 1 Supplementary Information

The following legend will be used throughout the supplemental proofs;

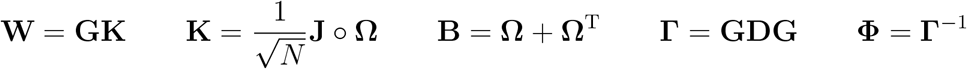

where **G** and **D** are diagonal matrices.

#### 1.1 General Form of Precision Matrix With Independent External Input

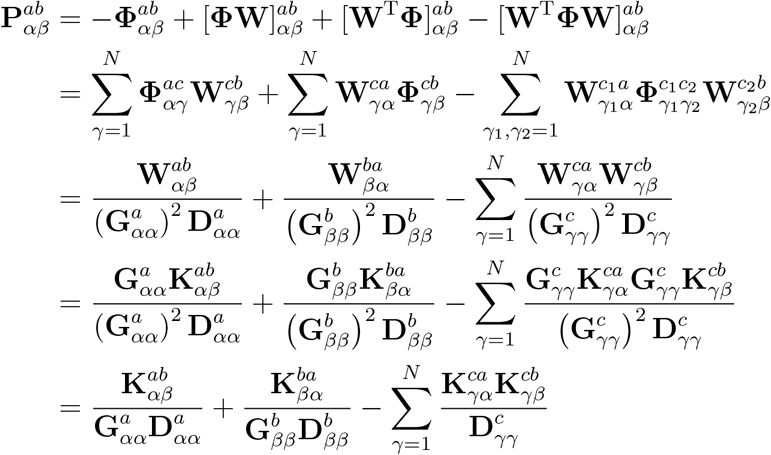

for *α* ≠ *β*.

#### 1.2 General Form of Discriminability of Precision Matrix With Independent External Input

##### 1.2.1 Signal

For general conditioning set **S**

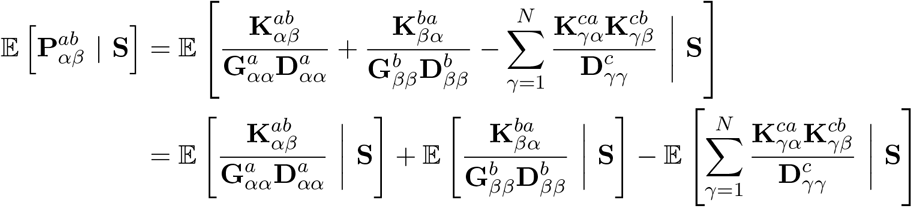

and since the main paper defines the conditioning sets in terms of mask *M*_1_(*a, b; m*), we implement this by taking **S** to correspond with 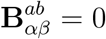 or 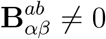. We begin by noting that for these sets, due to the fact that we have no self-loops (**W**_*αα*_ = 0 ∀*α*), we have the identity

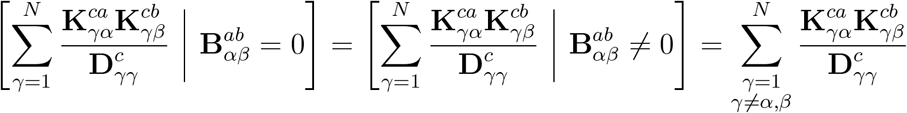

which directly implies that the signal (numerator of discriminability) of the precision in the case of independent external input reduces to

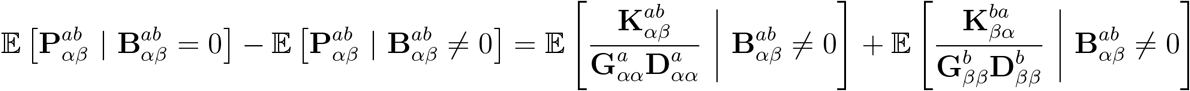

regardless of any specific structure (deterministic or stochastic) on **K** or **G**.

##### 1.2.2 Noise

Analogously,

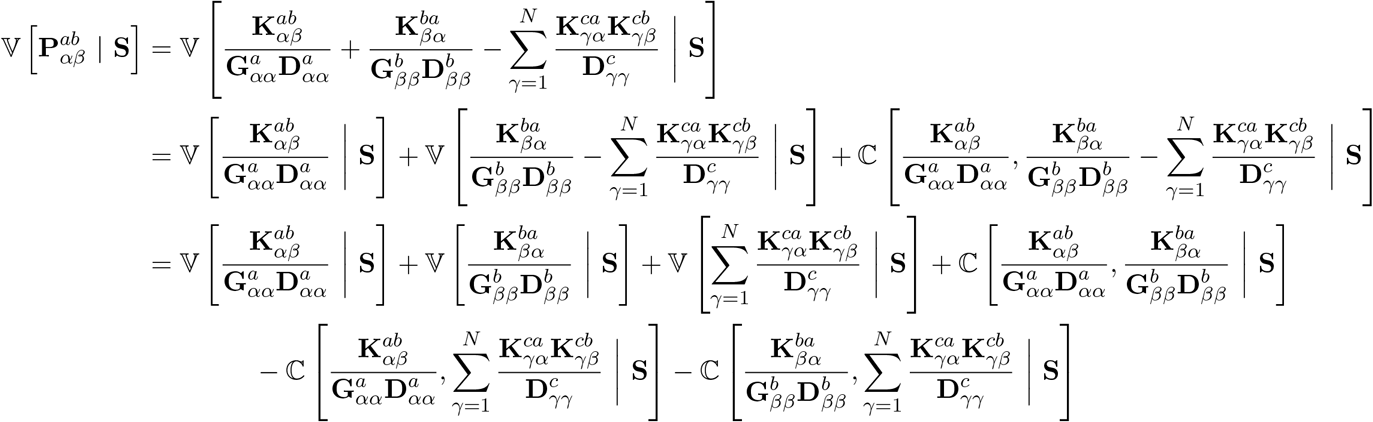

and applying the conditioning sets we find

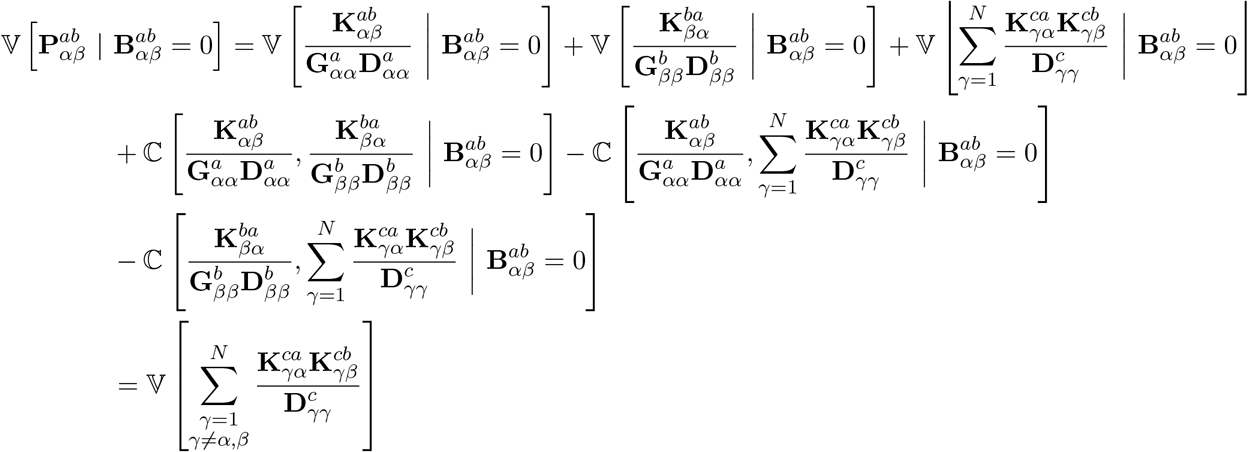

and

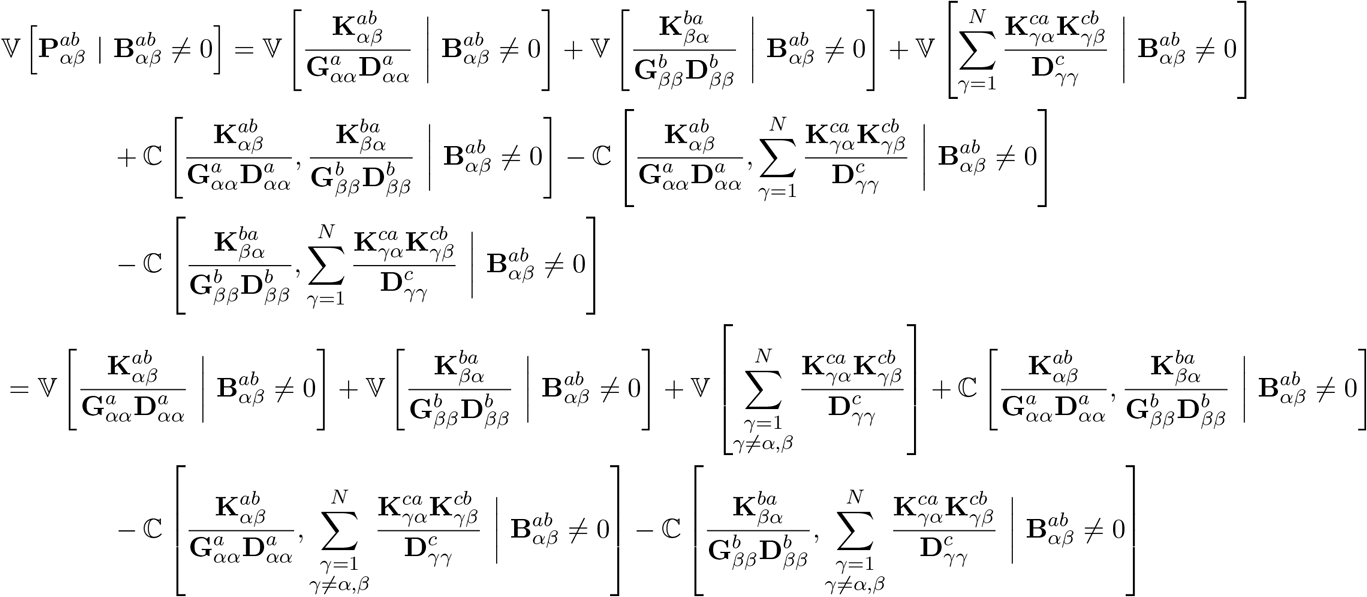

which implies the most general condition-specific noise form of

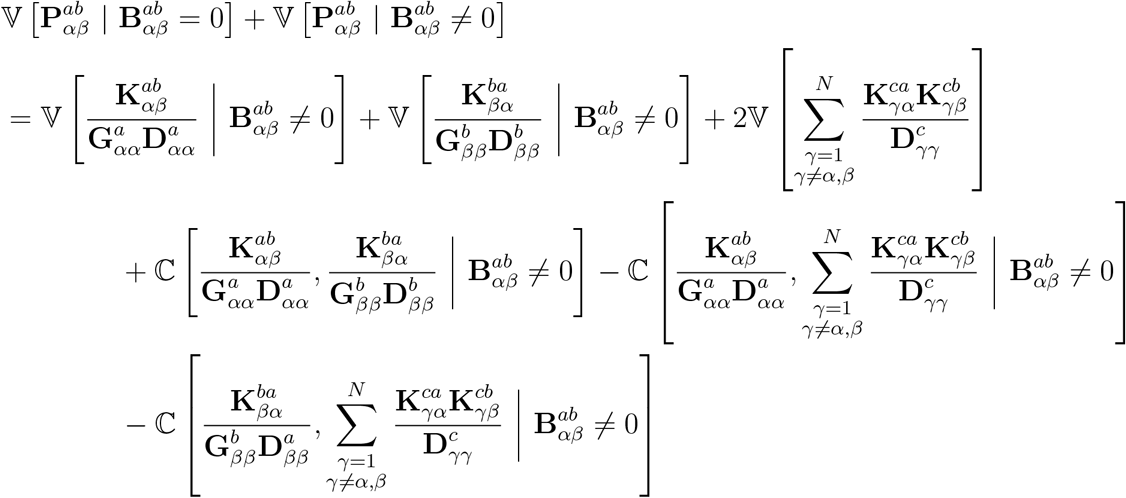

and since the signal is equivalent across all network structures, only the noise must be recalculated for each extended case.

#### 1.3 Discriminability of ER

Though specified in the main text, the following key will be helpful in understanding the simplification.

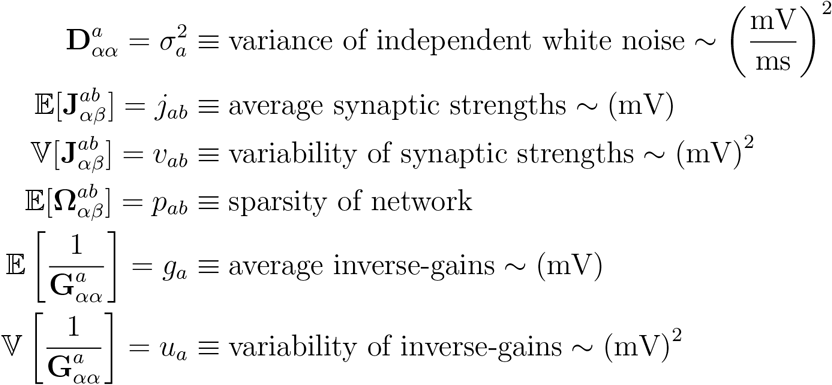

Assuming that 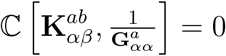 ∀*α, β*, the specific form of the signal for this case is

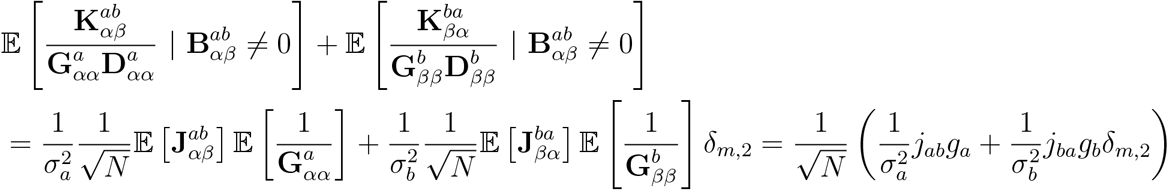

where it is important to note that since **B** is symmetric, specifying 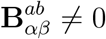 does not specify whether it is equal to 1 or 2 (a unidirectional or bidirectional motif) and so in simplifying discriminability *D* (*a, b; m*) for *m* = 1, 2 we must attach the *δ*_*m*,2_ on the transposed part of **W/K/J** following any evaluation of **B** ≠ 0. Since 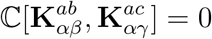 ∀*β* ≠ *γ*, the noise for the Erdos-Renyi case is

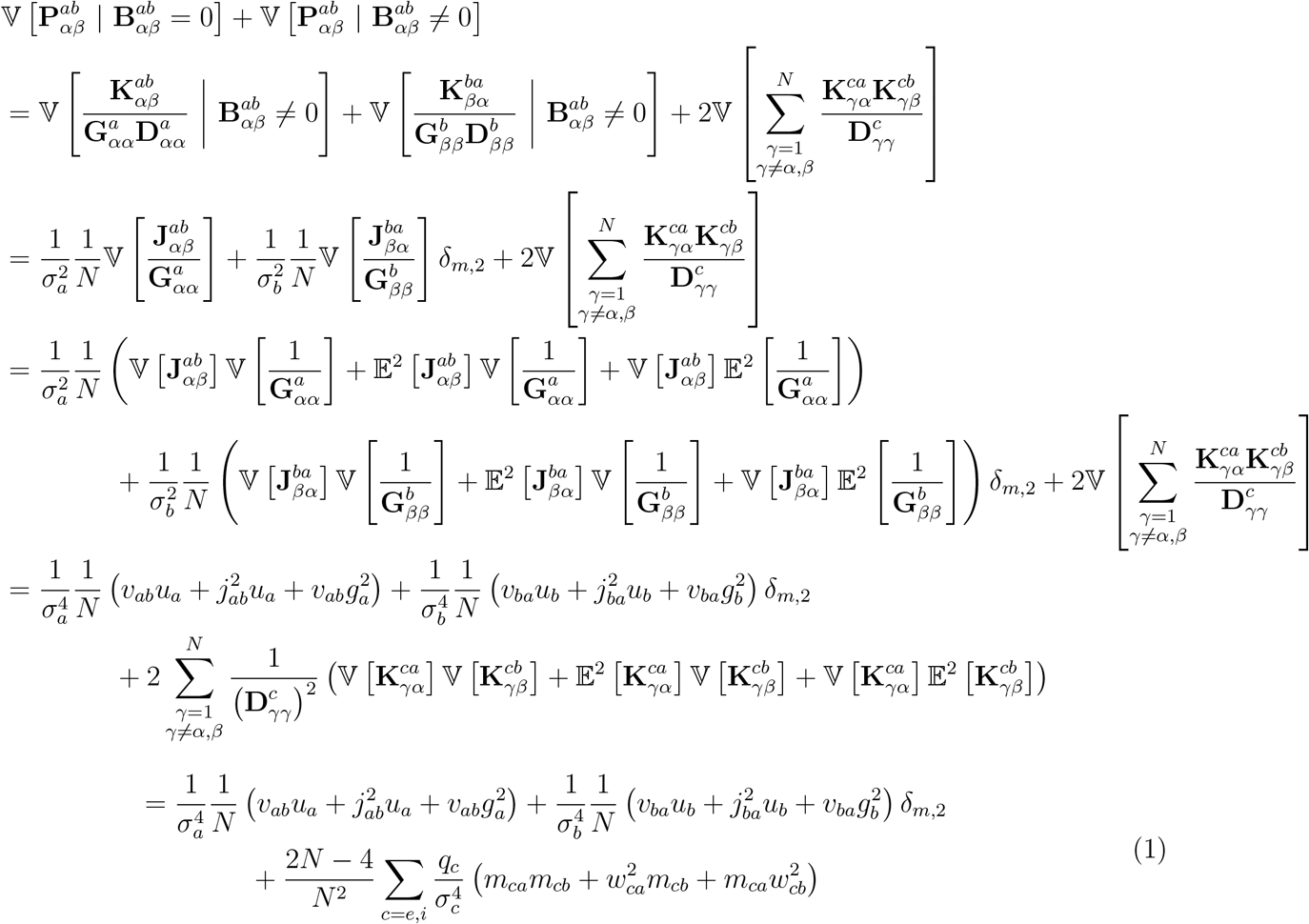

where

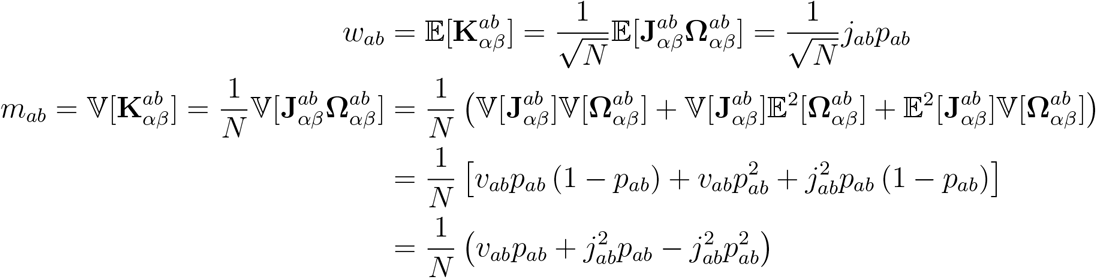

which together with the signal produces a final discriminability of

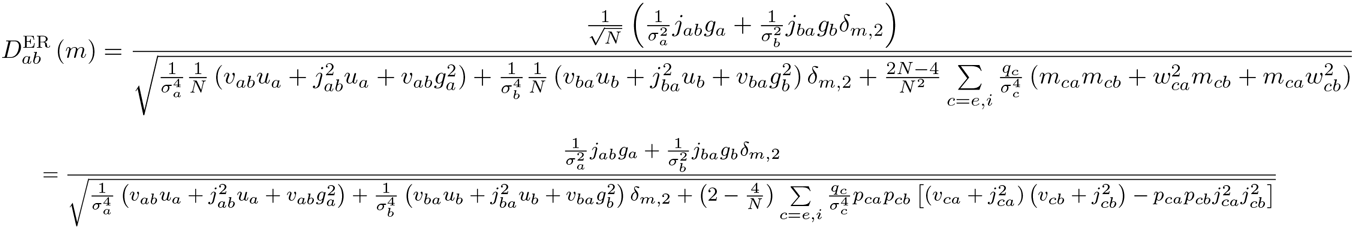

which can be simplified, assuming the gains and input noise are not population-specific (*i.e., g_a_* = *g, σ_a_ = σ*), as

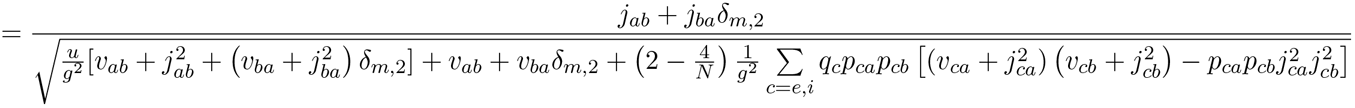

where the term 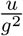 is the squared coefficient of variation (CV) of the gain distribution.

Note however that larger *v* causes slower convergence of the CLT by amplifying the variance of the distribution; *i.e*., there is less accuracy of the Gaussian assumption for the sum term at smaller *N* and thus can cause numerical differences between the theory and simulation. The Gaussian CLT outright fails to hold if *v* → ∞, though the resulting sums do converge to some α-stable distribution of unknown parameters and unknown corresponding analytical AUROC. Also note that in the case of an *α*-stable limit, likelihood ratio thresholding would have an even greater benefit to numerical inference. Larger *v* also induces greater truncation error on the synaptic distributions due to Dale’s Law, and thus even if the sum term is approximately Gaussian, the summation with the direct connectivity may cause some deviation from the theory. An alternative form of the equation using the alternate parameterization of

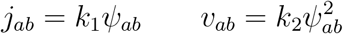

and if we take *g* = 1 and *u* = 0 as we do in the early-mid parts of the main paper, gives

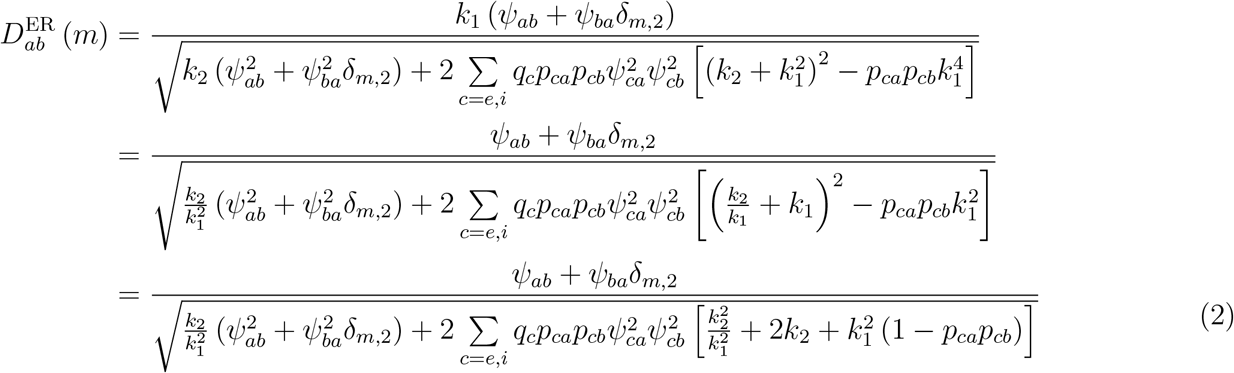

and this form will be slightly more useful for analyzing qualitative properties in Section 1.4.

##### 1.3.1 Discriminability of CER

Beyond this point, assume that *g_a_* = *g* =1 and *σ_a_* = *σ*. The resulting impact to the variability in **W** is

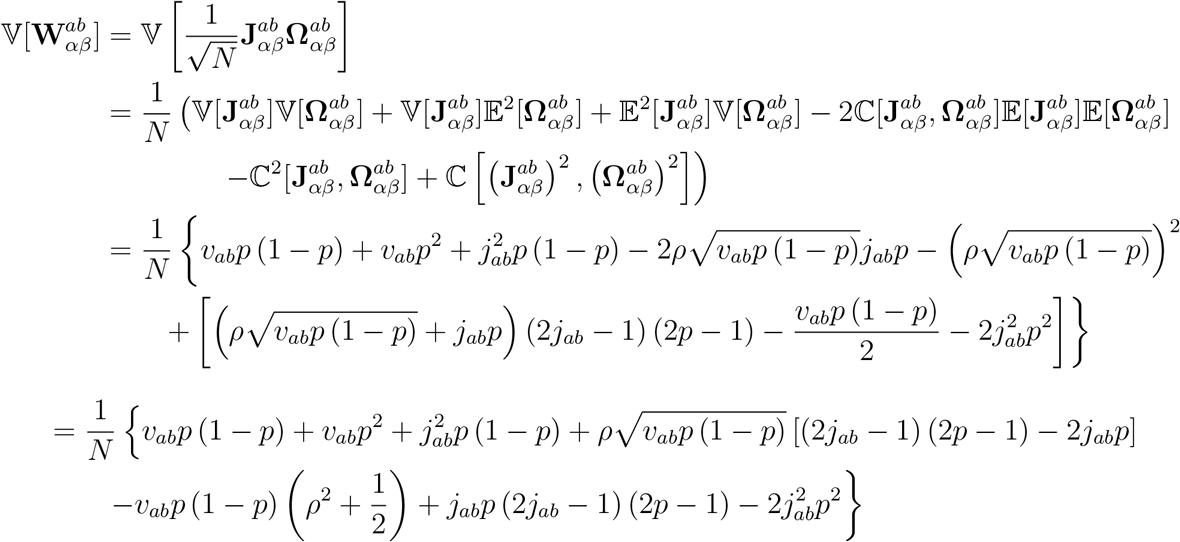

since

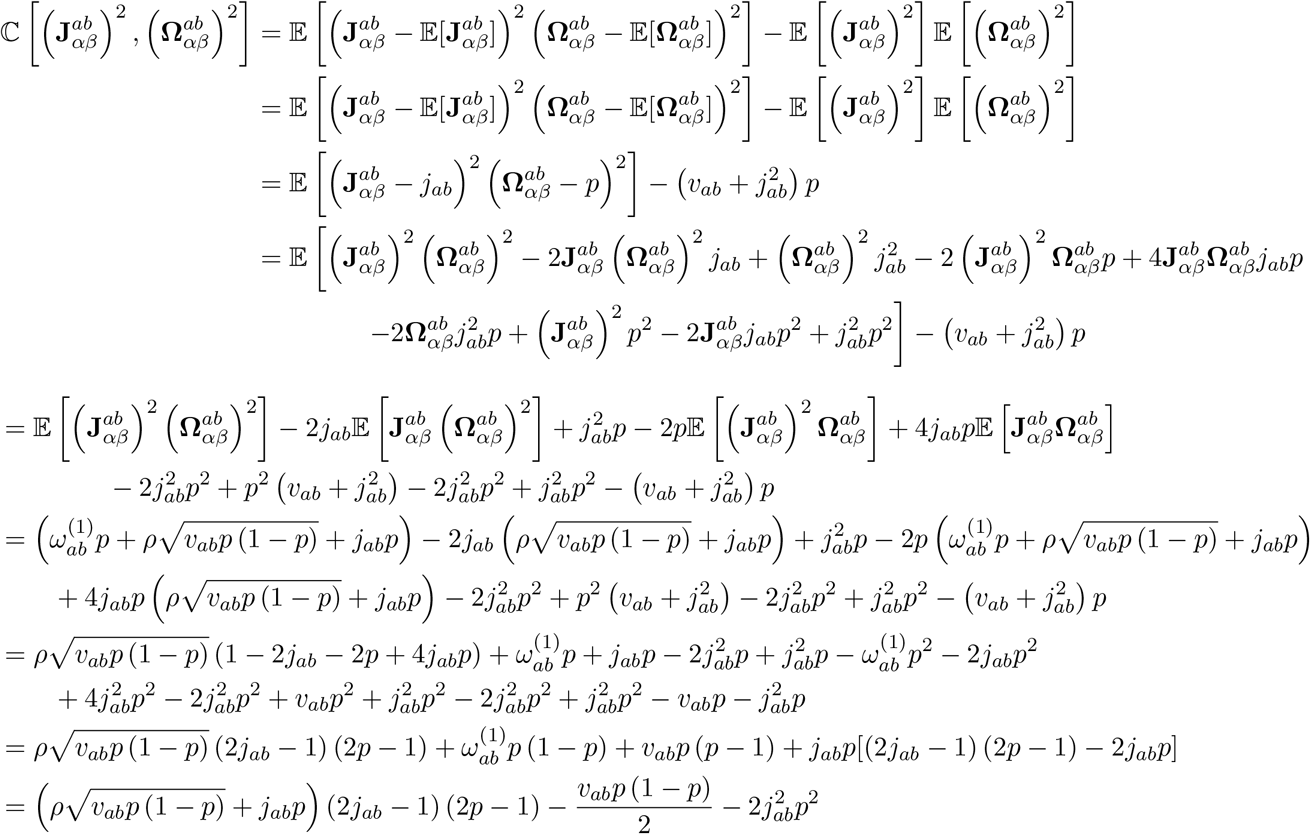

since

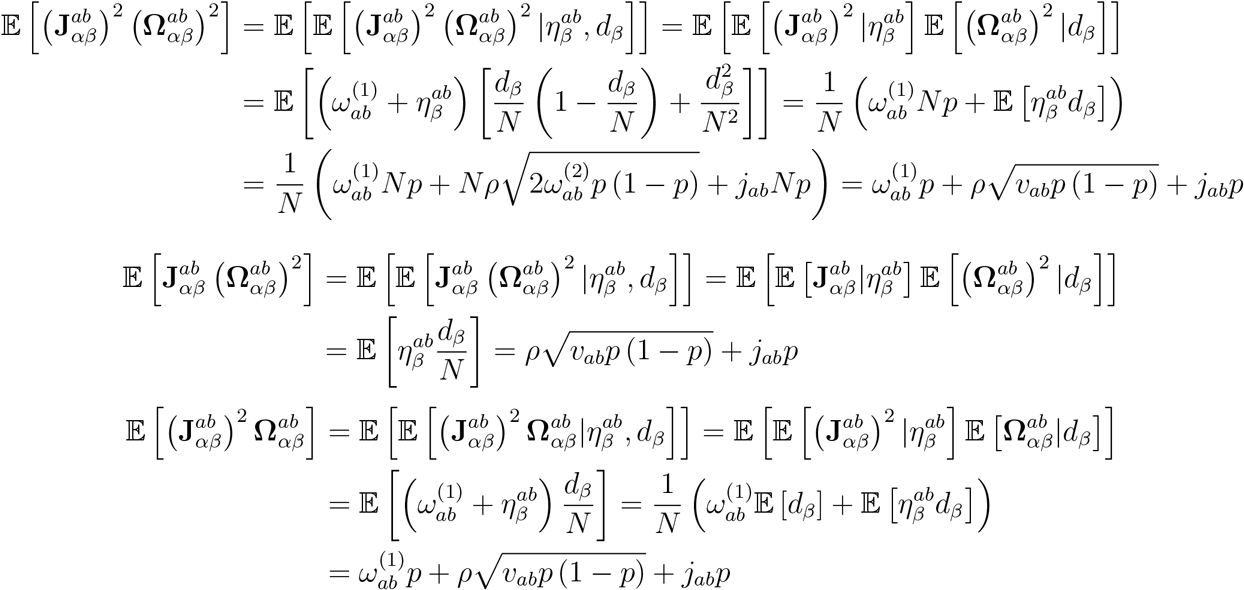

which finally allows for the identity

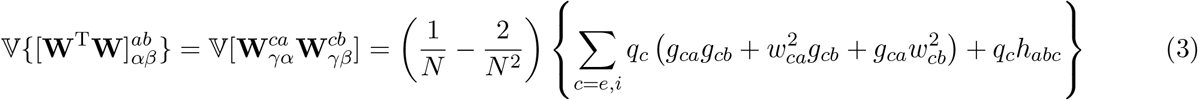

which implies

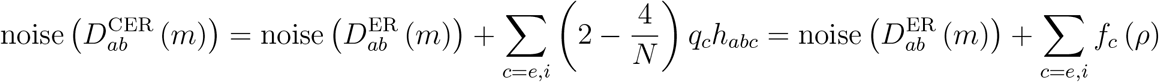

where

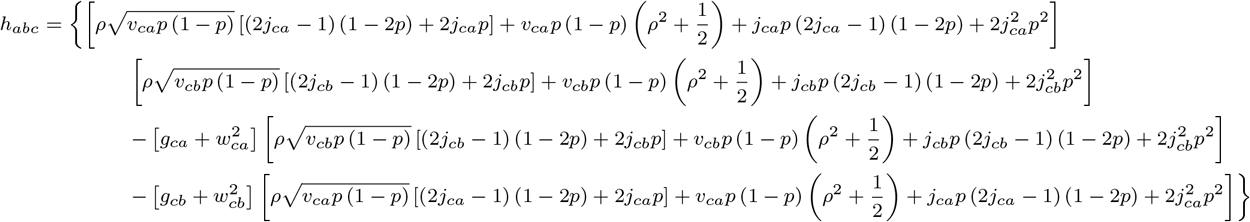

and though the sign of *h* can alternate depending on complicated relationships between the underlying variables, for our most common settings of 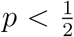 and 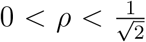 the sign is always positive ensuring that networks of this type have less discriminability than a corresponding Erdos-Renyi network.

##### 1.3.2 Discriminability of HD_in_

The most notable mathematical change for this model is to the variability of **Ω**, otherwise it follows the form of Eq. S.(1)

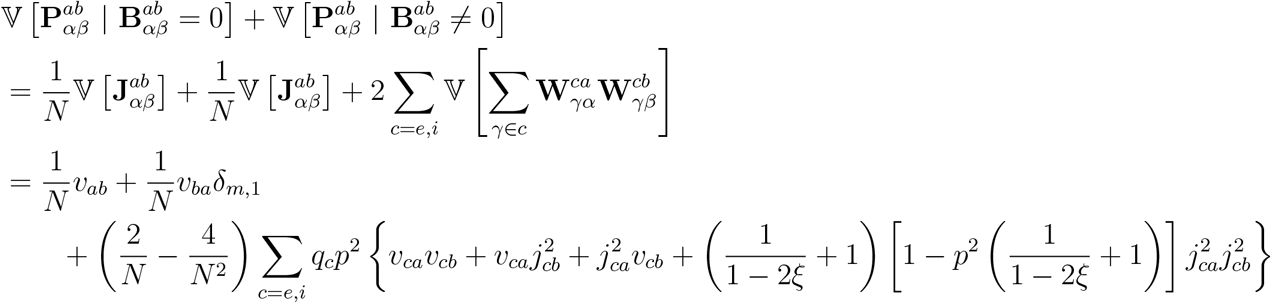

since

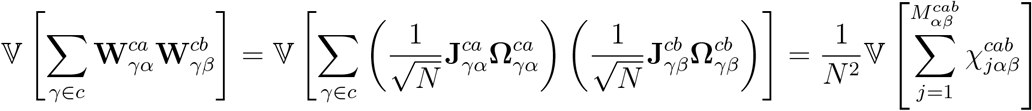

where *j* maps the non-zero indices of 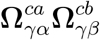, and where

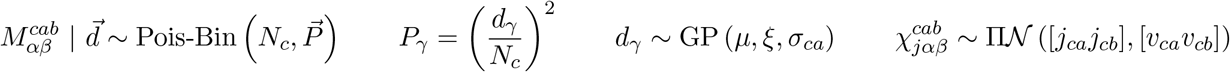

for ‘Pois-Bin’ indicating the Poisson-Binomial distribution, ‘GP’ denoting the Generalized Pareto distribution, and 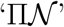 is the Product-Normal distribution. The additional variability from the random sum may be unconditionally expressed as

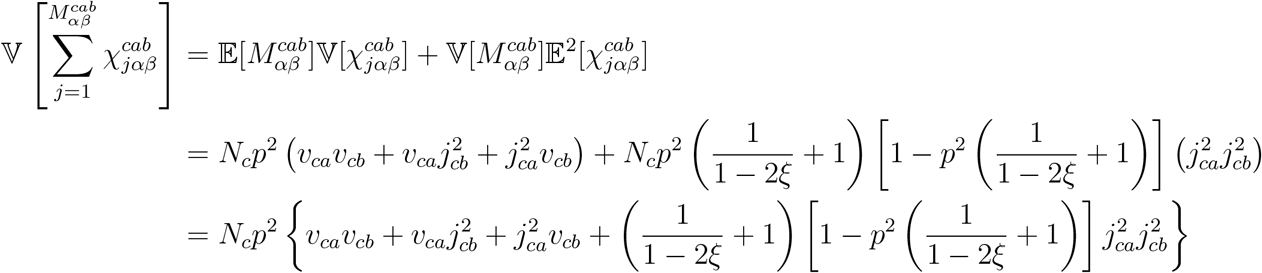

since

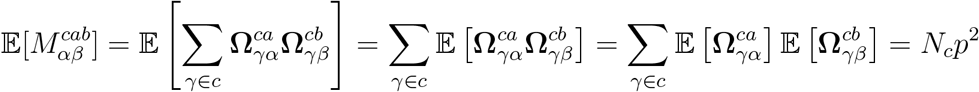

because

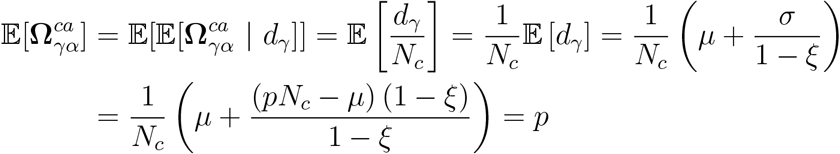

and then

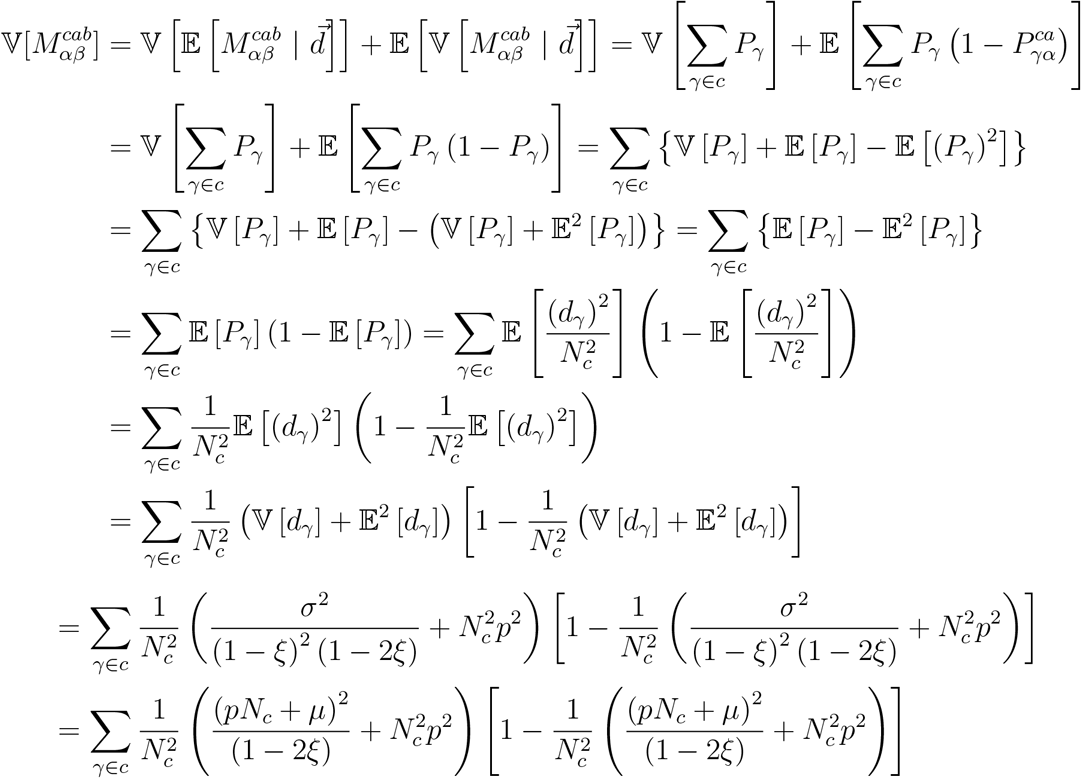

which for large *N* will be dominated by

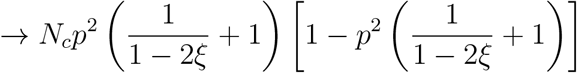

and combining all the relevant equations with simplification gives

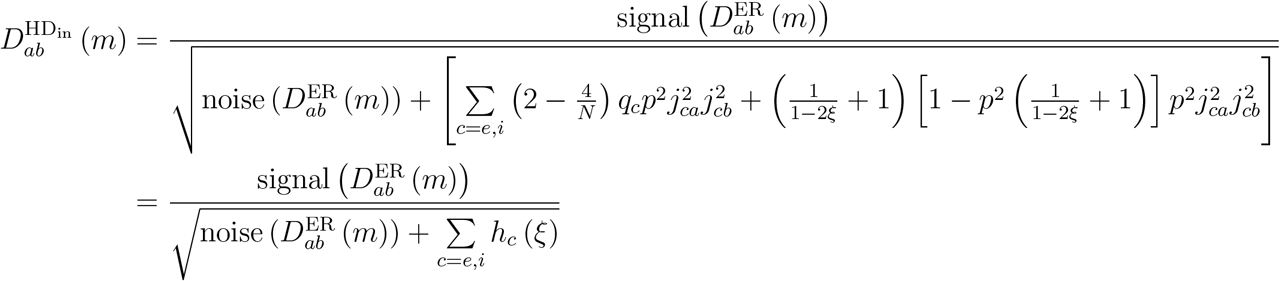

#### 1.4 Qualitative Properties

##### 1.4.1 Magnitude of Synaptic Strength

Using the alternative form of ER discriminability (Eq. S.(2)) it is easy to see that

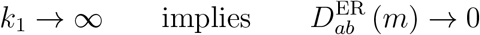

with more complex behavior occurring for smaller *k*_1_ values and two cases depending on *k*_2_. Optimizing over *k*_1_ holding all else fixed with *k*_2_ > 0, we find that *k*_1_ will have a maximum at

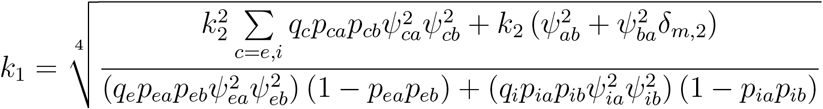

where it is crucial to note that if *k*_2_ = 0, then the maximum occurs as *k*_1_ → 0 which also leads to 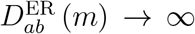. Otherwise, for *k*_2_ > 0, discriminability is no longer monotonic and will reach a maximum at the above value while going to zero for both *k*_1_ → 0 and *k*_1_ → ∞.

##### 1.4.2 Sparsity

Eq. S.(2) with global sparsity *p_ab_* = *p* is

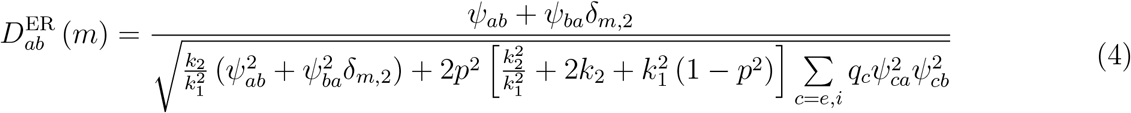

which, by holding everything beside *p* constant, can be viewed as

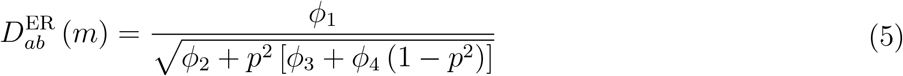

with all constants in 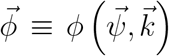 functionally dependent on one another and so cannot be altered independently. The constant resulting from the sum over *c* is absorbed into the inner constants of *ϕ*_3_,*ϕ*_4_. An applet that is useful in exploring this function is found in the accompanying code files; some numerical results indicate that the ratio of *k*_2_ to *k*_1_ determines the existence of a minima or monotonicity of the function, and that for certain blocks one is able to get non-monotonic behavior in changing *k*_1_ as well.

If we relax the global sparsity to being cell-type dependent on the presynaptic cell, that is *p_ab_* = *p_b_*, then we can analytically describe the minima as occurring along the curve

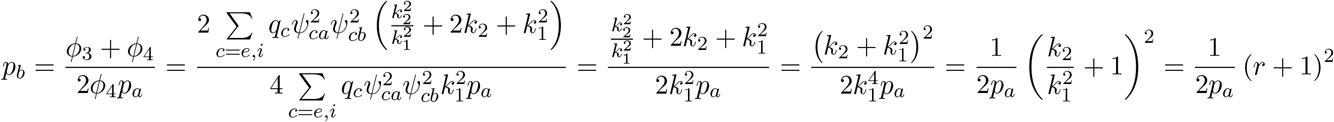

so long as *k*_2_ ≠ 0 (more precisely 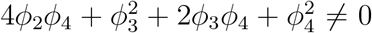) and that the equation is restricted to *p_b_* ∈ [0,1], else the minima occur outside the valid interval and the behavior appears monotonic over that range. Interestingly, this result theoretically holds no matter what the synaptic proportions are; it is only a function of the ratio *r* which is the magnitude of synaptic variability relative to the square of the magnitude of synaptic strength.

Further simplifying this analysis, if *p_a_ = p_b_ = p*, then it is simple to show that minima default to the right endpoint when *r* ≥ 1, and that the location of the minima prior to that vary linearly as 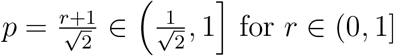 for *r* ∈ (0,1].

Alternatively, if *k*_2_ = 0 we get

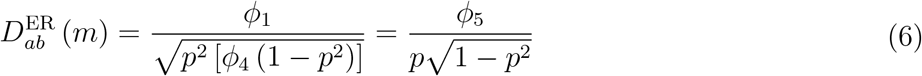

which has trivial minima fixed at 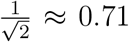 and so we maintain a smooth transition over the closed interval.

#### 1.5 Marginal Unbiased Estimate Discriminability - Samples vs. Optimal

Define

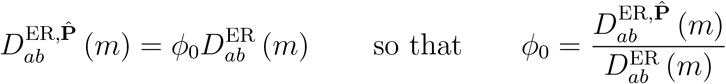

and by the previous assumption that the data powering the covariance estimation are Gaussian, that is

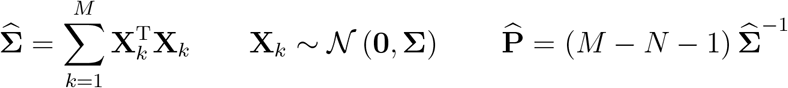

then the statistical estimator will follow an Inverse-Wishart distribution

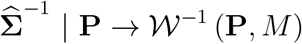

and then using the known variance of the Inverse-Wishart, the unconditional statistics are

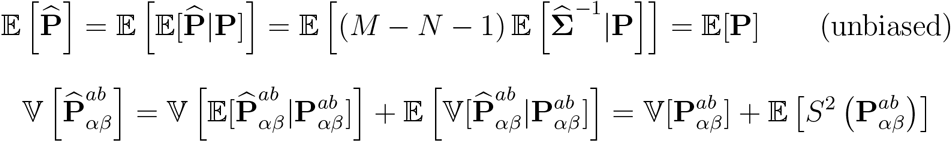

for the complicated conditional variance term 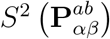 which also implies

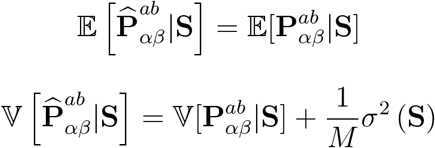

for any conditioning set **S** independent of the estimator. Here, *σ*^2^(**S**) is the unconditional average of *S*^2^(**P**|**S**) over **P**, and therefore only depends on the conditioning set. We have also here assumed that *M* is large enough such that the exact scaling approximates

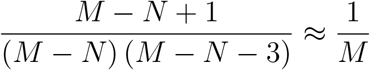

and altogether implies that

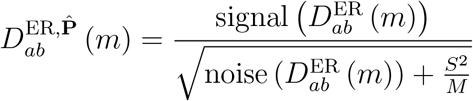

with 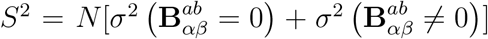 using our previous notation in dealing with **S** in terms of **B**. Combining this with *φ*_0_ gives the simplification of

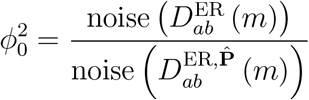

which may now be expanded as

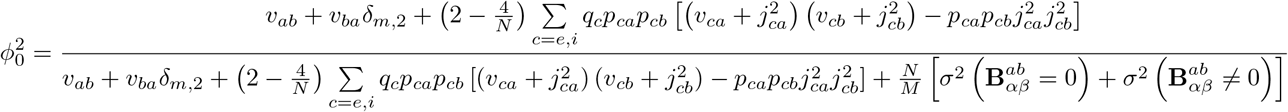

and can be solved for *M* as

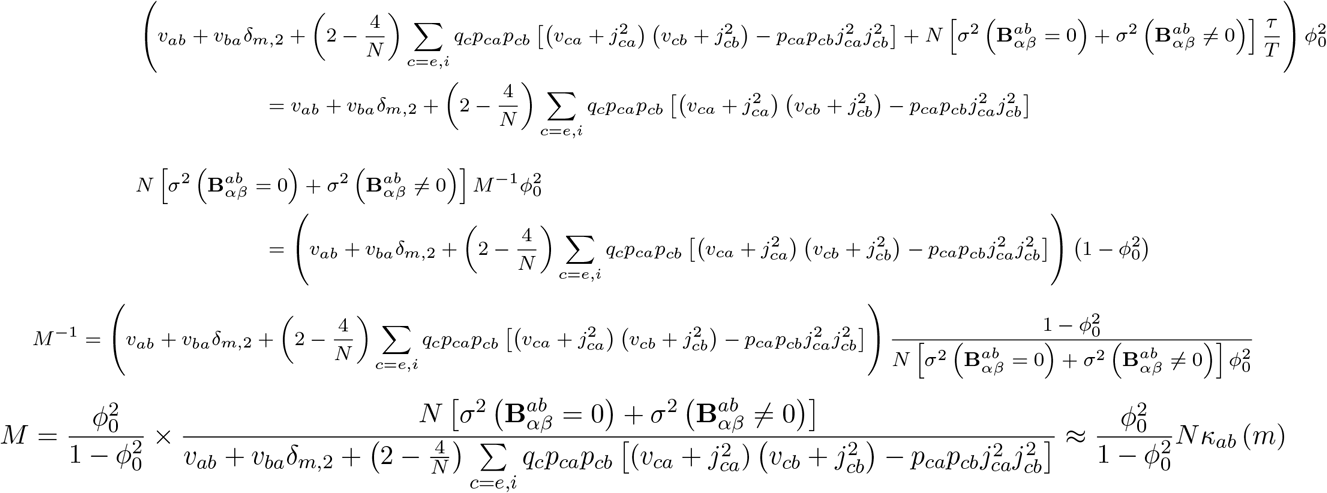

where the constant *κ_ab_*(*m*) is approximated as the 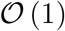 portion of the remaining terms with respect to *N*. This becomes particularly relevant to the scaling of *σ*^2^(**S**) with respect to *N*, which for different estimators may scale differently.

#### 1.6 Inverse Sample Covariance Estimate Discriminability

The previous section derives the general result for an arbitrary marginal estimate of precision. It should be noted that nearly all estimators of precision are taken as matrix functions, and marginal statistics are thus difficult to obtain. One of the only cases is for the inverse of sample covariance following an Inverse-Wishart distribution. In general, this is not recommended for any application; regularization methods such as GLASSO, sparse-latent methods, and many others are preferred to estimate global precision structure. However, all such methods are purely numerical - they have no known analytical form for their variability as estimators (in matrix or element-wise form) and this is why we use inverse sample covariance here. Intuitively, the variance of the numerical estimators ought to be smaller, and so this analysis serves as an upper bound on the amount of experiment time required to reach a desired percentage of the optimal discriminability. For the corresponding Inverse-Wishart, the rescaled marginal conditional variance are

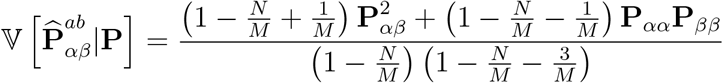

which under the large *M* setting required for the CLT to hold anyway, leads to the much simpler approximation of

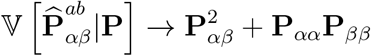

and so

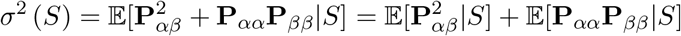

and it is not difficult to see that 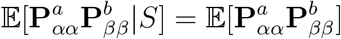 for our conditioning sets, we thus have only three more quantities we must compute. A more careful investigation reveals that

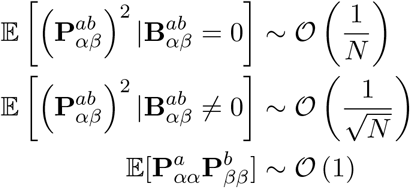

and so for the 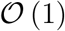 approximation to 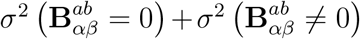 required for *κ* we need only the form of the third quantity. For this calculation, we must recall that from the general equation for precision there is an identity matrix that has often been dropped in the majority of special forms of precision throughout this paper as we have only previously been utilizing off-diagonal elements. Now it does adjust the diagonal elements as

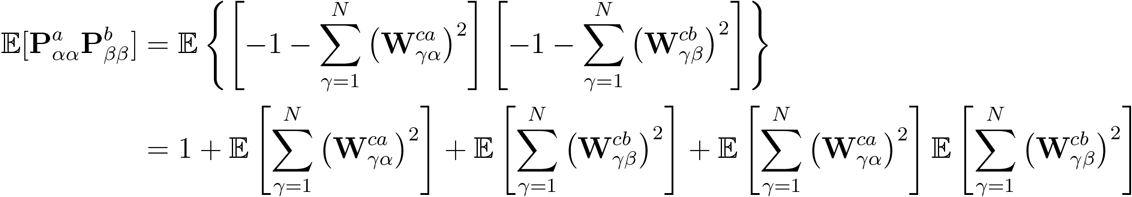

and since

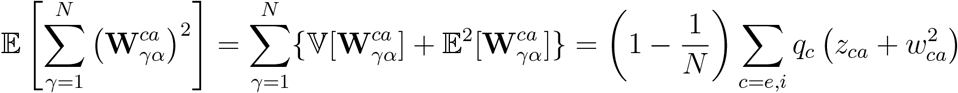

we may copy and adjust the indexing to get

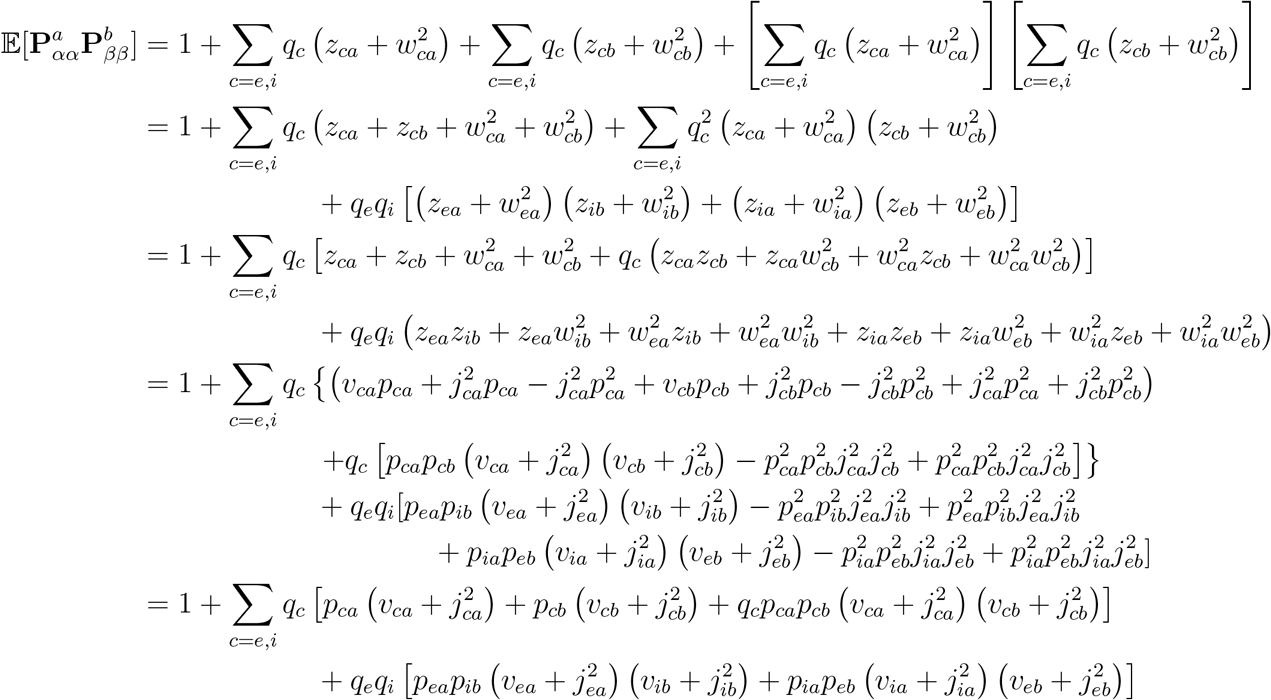

to largest order. Putting everything together, we find

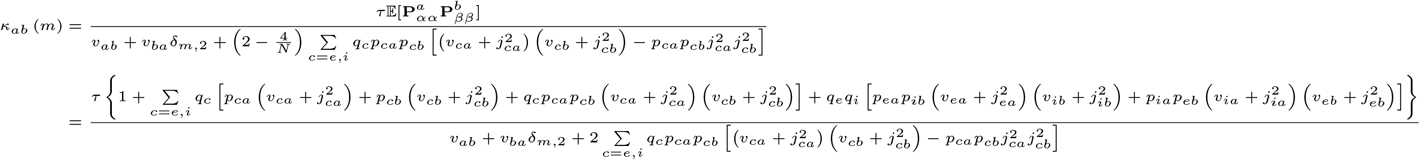

for this estimator.

#### 1.7 Supplementary Figures

**Figure S1:**
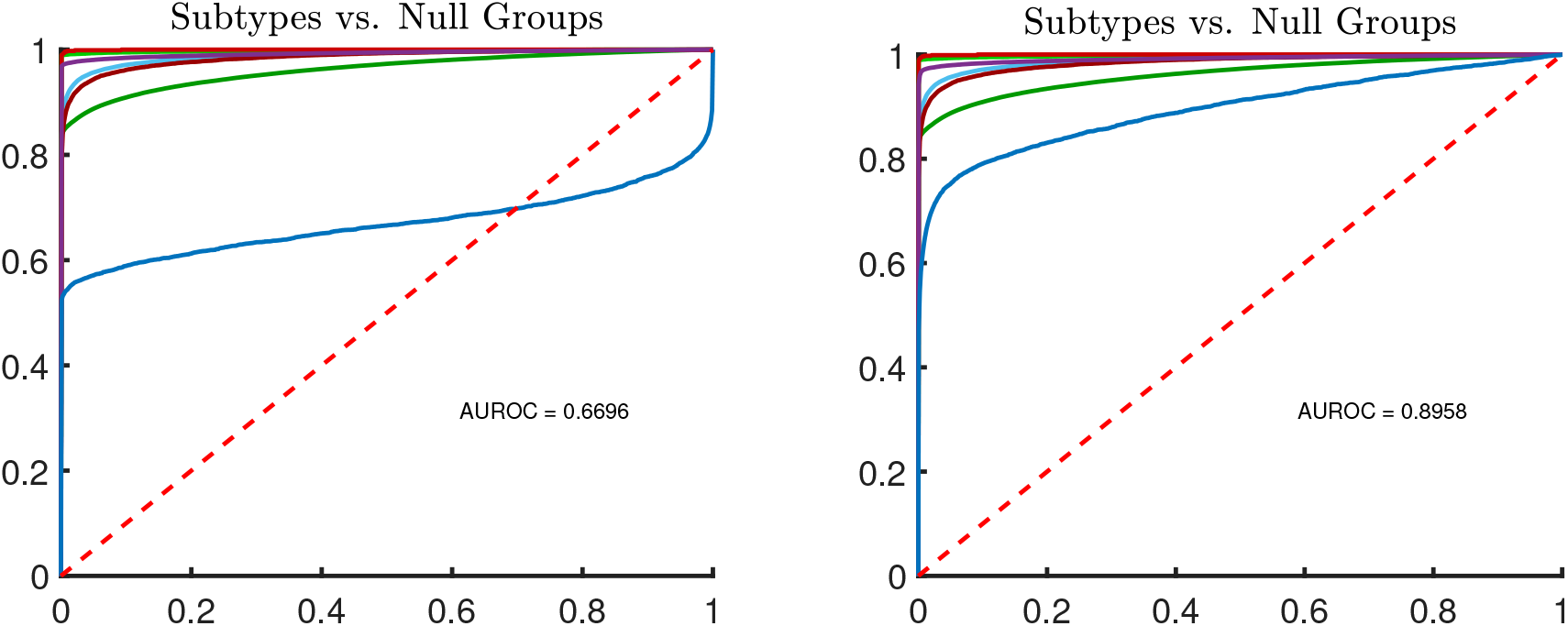
Example application of likelihood ratio thresholding improving inference whenever the original curves traversed the main diagonal. (a) ROC curves for low-strength, high-noise setting; same as Fig. 2h in the main text. (b) Same data, but scores are transformed according to the likelihood ratio.

**Figure S2:**
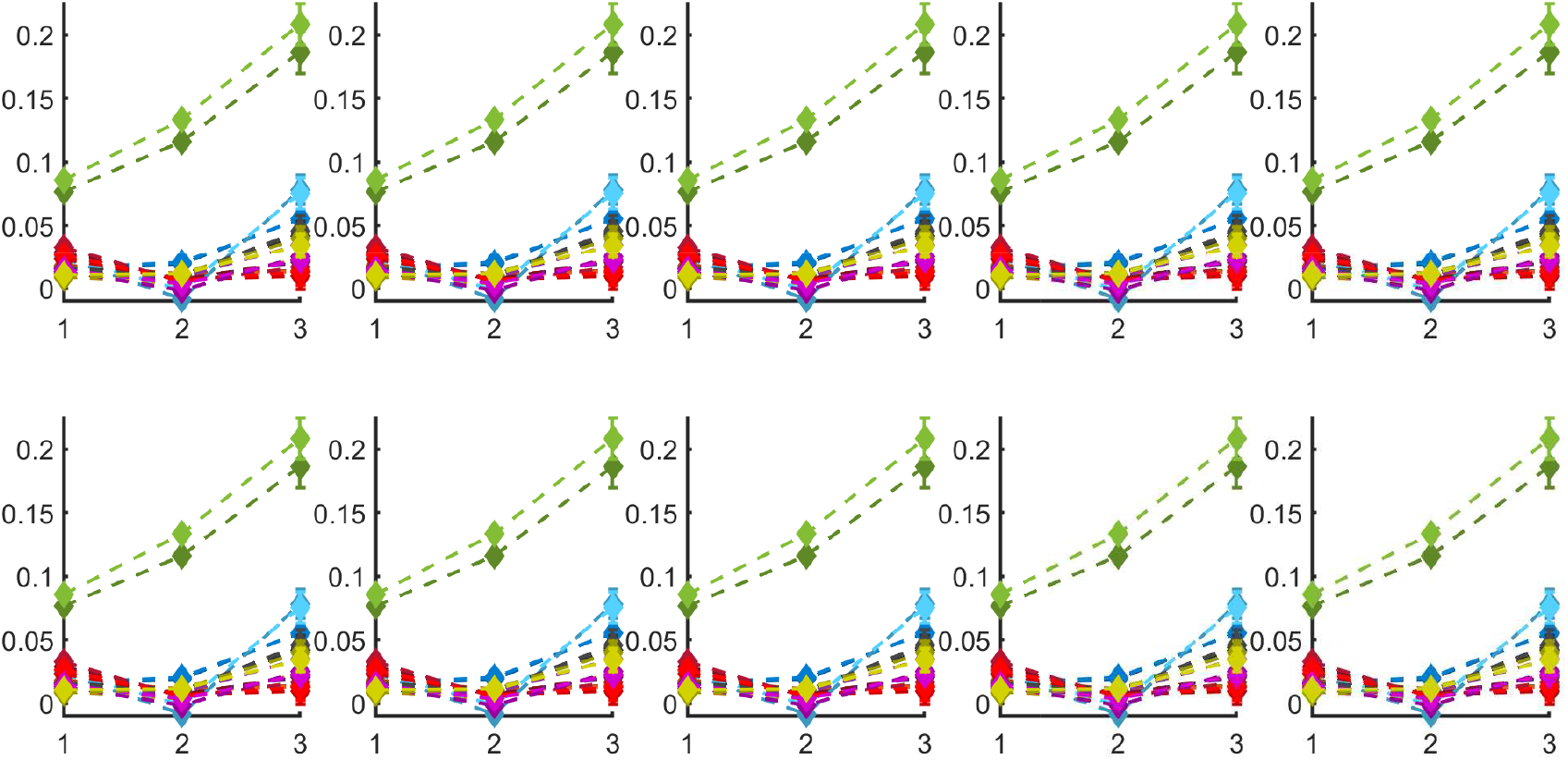
Same as Figures 8c from the main text, but over ten trials with randomly sampled hyperparameters.

**Figure S3:**
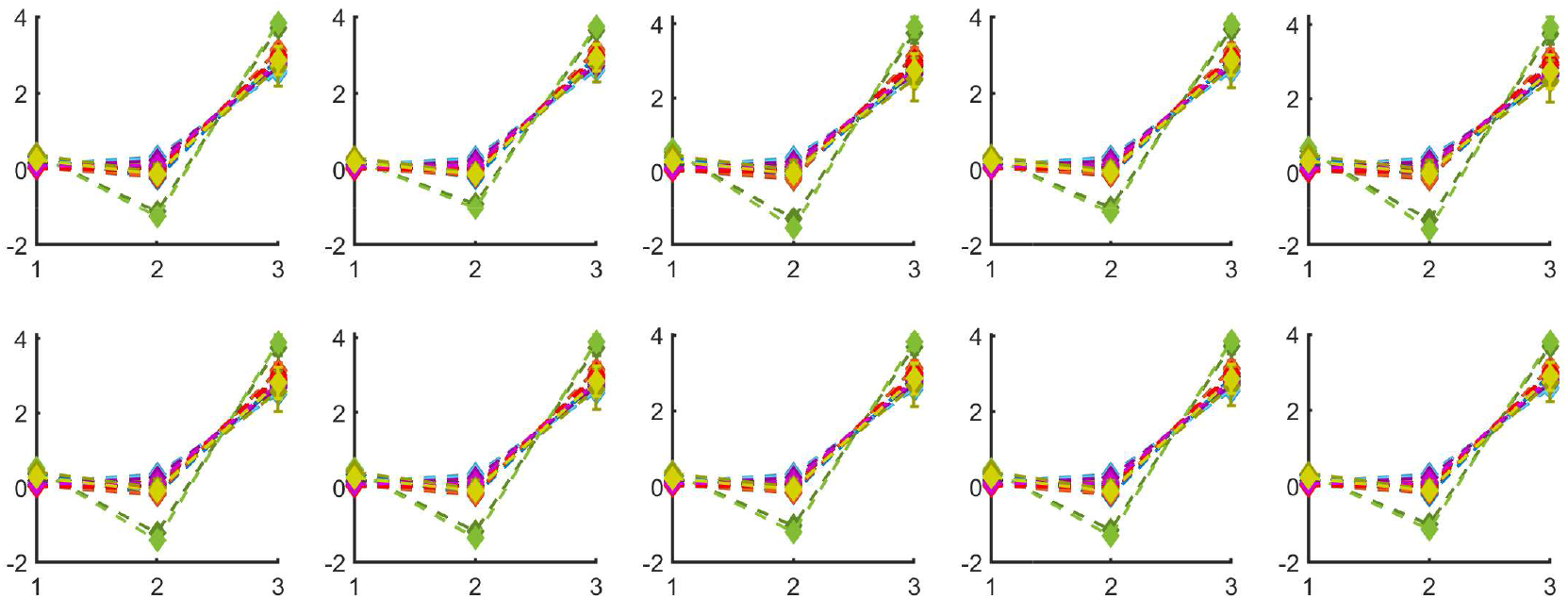
Same as Figures 8d from the main text, but over ten trials with randomly sampled hyperparameters.

**Figure S4:**
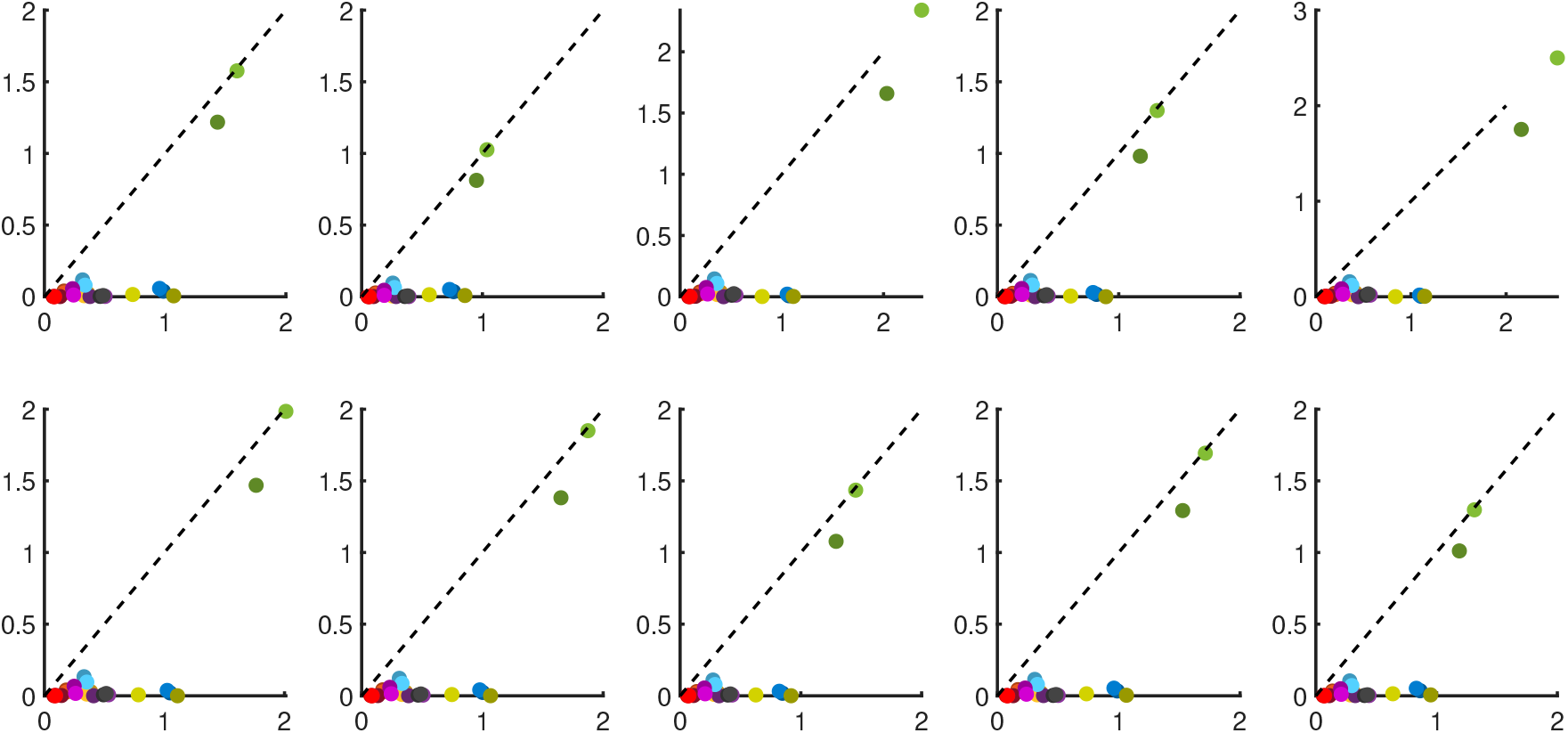
Same as Figures 8e from the main text, but over ten trials with randomly sampled hyperparameters.

**Figure S5:**
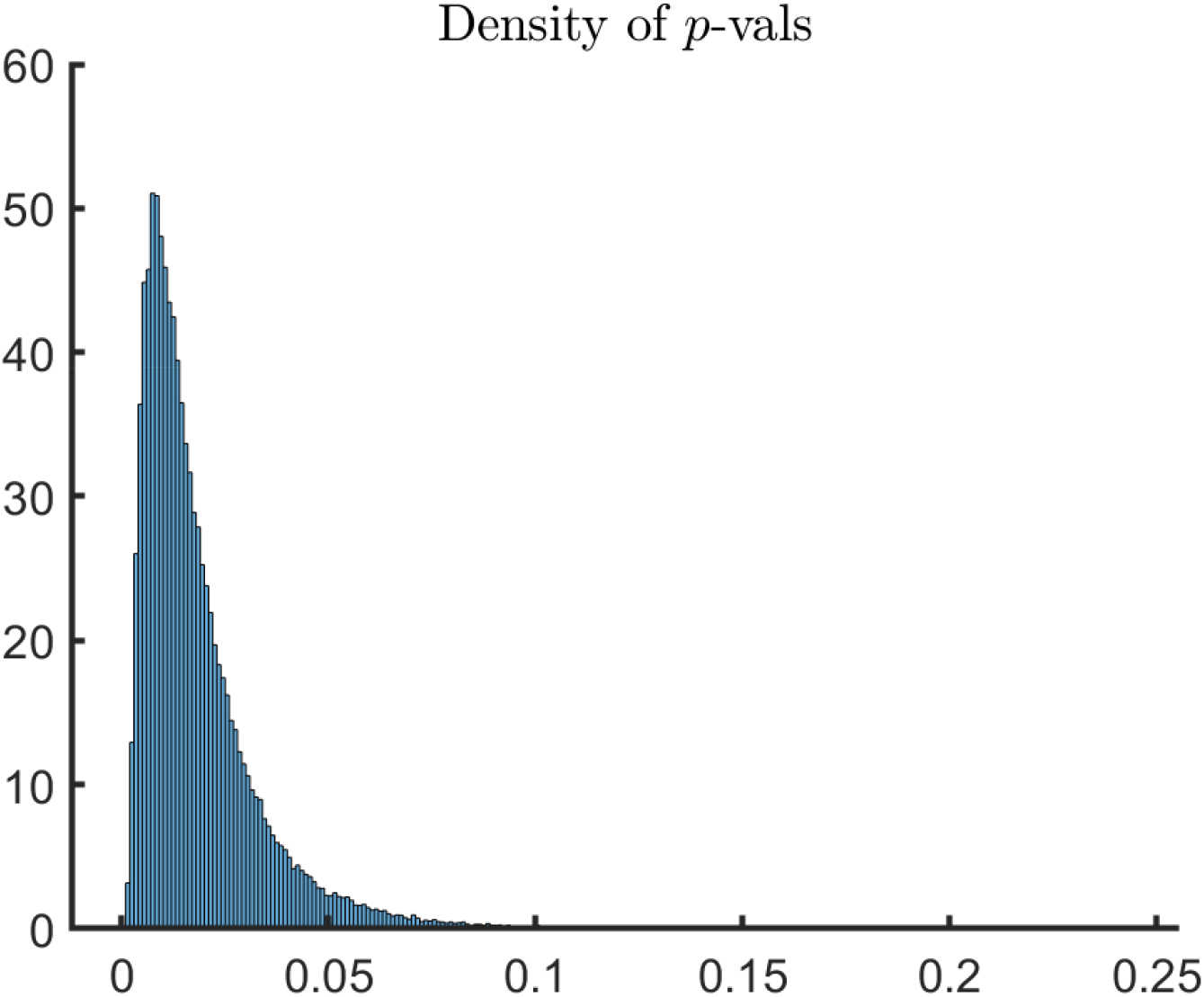
Distribution of *p*-values for the statistical test of mean-equality from section 11 of the main text using 100, 000 trials with randomly sampled hyperparameters. The mean is approximately 0.0185 with a maximum of 0.2421 and only 0.25% of the points greater than 0.1.

## References

Baker C, Ebsch C, Lampl I, Rosenbaum R (2018) The correlated state in balanced neuronal networks. bioRxiv p 372607

Barral J, D’Reyes A (2016) Synaptic scaling rule preserves excitatory-inhibitory balance and salient neuronal network dynamics. Nature Neuroscience 19(12):1690–1696

Bishop CM (2007) Pattern Recognition and Machine Learning

Brinkman BAW, Rieke F, Shea-Brown E, Buice MA (2017) Predicting how and when hidden neurons skew measured synaptic interactions pp 1–50

Chambers B, Levy M, Dechery1 JB, Maclean JN (2017) Ensemble stacking mitigates biases in inference of synaptic connectivity. Network Neuroscience Ensemble stacking mitigates biases in inference of synaptic connectivity. J N arXiv:1404.2263v1

Chiang AS, Lin CY, Chuang CC, Chang HM, Hsieh CH, Yeh CW, Shih CT, Wu JJ, Wang GT, Chen YC, Wu CC, Chen GY, Ching YT, Lee PC, Lin CY, Lin HH, Wu CC, Hsu HW, Huang YA, Chen JY, Chiang HJ, Lu CF, Ni RF, Yeh CY, Hwang JK (2011) Three-dimensional reconstruction of brain-wide wiring networks in drosophila at single-cell resolution. Current Biology 21(1):1–11

Cohen MR, Kohn A (2011) Measuring and interpreting neuronal correlations. Nature Neuroscience 14(7):811–819

Cotton RJ, Froudarakis E, Storer P, Saggau P, Tolias AS (2013) Three-dimensional mapping of microcircuit correlation structure. Frontiers in Neural Circuits

Dayan P, Abbott LF (2001) Theoretical neuroscience: computational and mathematical modeling of neural systems. MIT press

Doiron B, Litwin-Kumar A, Rosenbaum R, Ocker GK, JosiC K (2016) The mechanics of state-dependent neural correlations. Nature Neuroscience 19(3):383–393

Ebsch C, Rosenbaum R (2018) Imbalanced amplification: A mechanism of amplification and suppression from local imbalance of excitation and inhibition in cortical circuits. PLoS Computational Biology 14(3):1–28

Feldt S, Bonifazi P, Cossart R (2011) Dissecting functional connectivity of neuronal microcircuits: Experimental and theoretical insights

Friedrich J, Zhou P, Paninski L (2017) Fast online deconvolution of calcium imaging data. PLoS Computational Biology 1609.00639

Garaschuk O, Milos RI, Konnerth A (2006) Targeted bulk-loading of fluorescent indicators for two-photon brain imaging in vivo. Nature Protocols

Gardiner C (2009) Stochastic Methods - A Handbook for the Natural and Social Sciences. arXiv:1011.1669v3

Jiang X, Shen S, Cadwell CR, Berens P, Sinz F, Ecker AS, Patel S, Tolias AS (2016) Principles of connectivity among morphologically defined cell types in adult neocortex. Science 350(6264):1–21

Kadirvelu B, Hayashi Y, Nasuto SJ (2017) Inferring structural connectivity using Ising couplings in models of neuronal networks. Scientific Reports 7(1):1–12

Kalatsky VA, Stryker MP (2003) New paradigm for optical imaging: Temporally encoded maps of intrinsic signal. Neuron

Kohn A (2005) Stimulus Dependence of Neuronal Correlation in Primary Visual Cortex of the Macaque. Journal of Neuroscience

Krumin M, Reutsky I, Shoham S (2010) Correlation-Based Analysis and Generation of Multiple Spike Trains Using Hawkes Models with an Exogenous Input. Frontiers in Computational Neuroscience 4

Levy RB, Reyes AD (2012) in Mouse Primary Auditory Cortex 32(16):5609–5619

Lin TW, Das A, Krishnan GP, Bazhenov M, Sejnowski TJ (2017) Differential covariance: A new class of methods to estimate sparse connectivity from neural recordings. Neural Computation 29(10):2581–2632, 1706.02451

Lütcke H, Gerhard F, Zenke F, Gerstner W, Helmchen F (2013) Inference of neuronal network spike dynamics and topology from calcium imaging data. Frontiers in Neural Circuits 7(December):1–20

Magrans de Abril I, Yoshimoto J, Doya K (2018) Connectivity inference from neural recording data: Challenges, mathematical bases and research directions. Neural Networks 102:120–137, 1708.01888

Maswadeh WM, Snyder PS (2012) Multivariable and multigroup Receiver Operating Characteristics curve analyses for qualitative and quantitative analysis. Edgewood Chemical Biological Center ECBC-TR-92(U.S. Army Research, Development and Engineering Command)

Mishchencko Y, Vogelstein J, Paninski L (2007) a Bayesian Approach for Inferring Neuronal. Statistics arXiv:1107.4228v1

Paninski L (2004) Maximum likelihood estimation of cascade point-process neural encoding models. Network: Computation in Neural Systems

Pernice V, Rotter S (2013) Reconstruction of sparse connectivity in neural networks from spike train covariances. Journal of Statistical Mechanics: Theory and Experiment 2013(3)

Pernice V, Staude B, Cardanobile S, Rotter S (2011) How structure determines correlations in neuronal networks. PLoS Computational Biology 7(5)

Pfeffer CK, Xue M, He M, Huang ZJ, Scanziani M (2013) Inhibition of inhibition in visual cortex: The logic of connections between molecularly distinct interneurons. Nature Neuroscience DOI 10.1038/nn.3446, NIHMS150003

Pillow JW, Shlens J, Paninski L, Sher A, Litke AM, Chichilnisky EJ, Simoncelli EP (2008) Spatio-temporal correlations and visual signalling in a complete neuronal population. Nature NIHMS150003

Pnevmatikakis EA, Soudry D, Gao Y, Machado TA, Merel J, Pfau D, Reardon T, Mu Y, Lacefield C, Yang W, Ahrens M, Bruno R, Jessell TM, Peterka DS, Yuste R (2017) HHS Public Access 89(2):285–299

Poli D, Pastore VP, Martinoia S, Massobrio P (2016) From functional to structural connectivity using partial correlation in neuronal assemblies. Journal of Neural Engineering 13(2):26,023

Pyle R, Rosenbaum R (2016) Highly connected neurons spike less frequently in balanced networks. Physical Review E 93(4):1–6, 1601.04972

Renart A, Rocha JD, Bartho P, Hollender L, Reyes A, Harris KD (2010) The Asynchronus State in Cortical Circuits. Science 327(5965):587–590, 1002.1037

Rosenbaum R, Smith MA, Kohn A, Rubin JE, Doiron B (2017) The spatial structure of correlated neuronal variability. Nature Neuroscience 20(1):107–114

Singh R, Ghosh D, Adhikari R (2017) Fast Bayesian inference of the multivariate Ornstein-Uhlenbeck process 012136:1–9, 1706.04961

Smith MA, Kohn A (2008) Spatial and Temporal Scales of Neuronal Correlation in Primary Visual Cortex. Journal of Neuroscience

Song S, Sjöström PJ, Reigl M, Nelson S, Chklovskii DB (2005) Highly nonrandom features of synaptic connectivity in local cortical circuits. In: PLoS Biology, DOI 10.1371/journal.pbio.0030068

Soudry D, Keshri S, Stinson P, Oh Mh, Iyengar G, Paninski L (2013) A shotgun sampling solution for the common input problem in neural connectivity inference pp 1–10, 1309.3724

Trousdale J, Hu Y, Shea-Brown E, Josić K (2012) Impact of network structure and cellular response on spike time correlations. PLoS Computational Biology 8(3), 1110.4914

Van Vreeswijk C, Sompolinsky H (1996) Chaos in neuronal networks with balanced excitatory and inhibitory activity. Science 274(5293):1724–1726, DOI 10.1126/science.274.5293.1724, 1011.1669

Vinci G, Smith MA, Kass RE (2018) Adjusted regularization of cortical covariance. Journal of Computational Neuroscience DOI 10.1007/s10827-018-0692-x

Vogelstein JT, Packer AM, Machado TA, Sippy T, Yuste R, Paninski L, Babadi B (2012) Fast Nonnegative Deconvolution for Spike Train Inference From Population Calcium Imaging Fast Nonnegative Deconvolution for Spike Train Inference From Population Calcium Imaging. Journal of neurophysiology

van Vreeswijk C, Sompolinsky H (1998) Chaotic Balanced State in a Model of Cortical Circuits. Neural Computation 10(6):1321–1371, arXiv:1011.1669v3

Widloski J, Marder MP, Fiete IR (2018) Inferring circuit mechanisms from sparse neural recording and global perturbation in grid cells. eLife DOI 10.7554/eLife.33503

Yaglom A (1962) An Introduction to the Theory of Stationary Random Functions

Yatsenko D, Josić K, Ecker AS, Froudarakis E, Cotton RJ, Tolias AS (2015) Improved Estimation and Interpretation of Correlations in Neural Circuits. PLoS Computational Biology 11(3): 1–28

Yatsenko D, Froudarakis E, Ecker A, Rosenbaum R, Josić K, Tolias A (2016) Strong functional connectivity of parvalbumin-expressing cortical interneurons. Computational and Systems Neuroscience Meeting (COSYNE 2016)

Zaytsev YV, Morrison A, Deger M (2015) Reconstruction of recurrent synaptic connectivity of thousands of neurons from simulated spiking activity. Journal of Computational Neuroscience 39(1):77–103, 1502.04993

